# Dissociable effects of visual crowding on the perception of colour and motion

**DOI:** 10.1101/639450

**Authors:** John A. Greenwood, Michael J. Parsons

## Abstract

Our ability to recognise objects in peripheral vision is fundamentally limited by crowding, the deleterious effect of clutter that disrupts the recognition of features ranging from orientation and colour to motion and depth. Prior research is equivocal on whether this reflects a singular process that disrupts all features simultaneously or multiple processes that affect each independently. We examined crowding for motion and colour, two features that allow a strong test of feature independence. ‘Cowhide’ stimuli were presented 15 degrees in peripheral vision, either in isolation or surrounded by flankers to give crowding. Observers reported either the target direction (clockwise/counterclockwise from upwards) or its hue (blue/purple). We first established that both features show systematic crowded errors (predominantly biased towards the flanker identities) and selectivity for target-flanker similarity (with reduced crowding for dissimilar target/flanker elements). The multiplicity of crowding was then tested with observers identifying both features: a singular object-selective mechanism predicts that when crowding is weak for one feature and strong for the other that crowding should be all-or-none for both. In contrast, when crowding was weak for colour and strong for motion, errors were reduced for colour but remained for motion, and vice versa with weak motion and strong colour crowding. This double dissociation reveals that crowding disrupts certain combinations of visual features in a feature-specific manner, ruling out a singular object-selective mechanism. The ability to recognise one aspect of a cluttered scene, like colour, thus offers no guarantees for the correct recognition of other aspects, like motion.

**Significance statement:** Our peripheral vision is primarily limited by crowding, the disruption to object recognition that arises in clutter. Crowding is widely assumed to be a singular process, affecting all of the features (orientation, motion, colour, etc.) within an object simultaneously. In contrast, we observe a double dissociation whereby observers make errors regarding the colour of a crowded object whilst correctly judging its direction, and vice versa. This dissociation can be reproduced by a population-coding model where the direction and hue of target/flanker elements are pooled independently. The selective disruption of some object features independently of others rules out a singular crowding mechanism, posing problems for high-level crowding theories, and suggesting that the underlying mechanisms may be distributed throughout the visual system.

## Introduction

Our ability to recognise objects declines sharply in peripheral vision (1). This is not simply due to resolution or acuity – objects that are visible in isolation become indistinguishable when other objects fall within surrounding ‘interference zones’ (2–4). This process, known as crowding, presents the fundamental limit on peripheral vision, with pronounced elevations in central vision in disorders including amblyopia (5) and dementia (6).

Crowded impairments arise due to a systematic change in the appearance of target objects (7, 8), particularly outside the fovea (9) where targets are induced to appear more similar to nearby ‘flankers’. Crowding disrupts the recognition of features throughout the visual system, including orientation (10), position (11), colour (12, 13), motion (14), and depth (15). Within these dimensions, crowding is also modulated by the similarity between target/flanker elements – differences in features including orientation and colour reduce errors considerably (10, 16). Given the distributed processing of these features across the visual system (17, 18), can one process produce this multitude of effects? Most models implicitly assume that crowding is a single mechanism that affects all features in a combined manner, particularly for higher-order approaches where crowding derives from attention (19, 20) or grouping (21). If crowding were instead to operate independently for distinct visual features, these effects could involve an array of neural substrates with varied mechanisms.

A key prediction for a combined crowding process is that a release from crowding in one feature domain (e.g. colour) should release other features (e.g. motion) at the same time. Accordingly, target-flanker differences in colour or contrast polarity can reduce crowding for judgements of spatial form (16), while differences in orientation improve crowded position judgements (22). Others have however found that judgements of spatial frequency, colour, and orientation show a mixture of independent and combined errors (23). This discrepancy may reflect the specific features used in each study. Here we examined whether crowding is combined or independent for judgements of motion and colour – arguably the two features with the clearest separation in the visual system (17, 18).

We conducted 3 experiments with motion and colour, each using cowhide-like stimuli (24, 25) in the upper visual field. Experiments 1 and 2 examined crowding for each feature separately to determine both the nature of the errors (i.e. their systematicity) and the flanker conditions that give strong vs. weak crowding. We then measured the independence of crowding with conjoint motion/colour judgements in Experiment 3 by selecting conditions where crowding was strong for one feature and weak for the other, or vice versa.

## Results

In Experiment 1, observers viewed moving cowhide stimuli and reported the movement direction (clockwise/counterclockwise of upwards) of a target presented either in isolation or surrounded by flankers moving in 1 of 16 directions (Figure 1A; Movie S1). Example data are shown in Figure 1B, where unflanked judgments (grey points) transition rapidly from predominantly CW to CCW at directions around upwards (0°). The psychometric function accordingly shows low bias in the point of subjective equality with upwards (PSE; the 50% midpoint), with the steep slope indicating a low threshold (the difference from 50% to 75% CCW responses). With upwards-moving flankers (+0°; blue points), performance declined, with a shallower psychometric function, but nonetheless remained unbiased. In contrast, flankers moving 30° CCW of upwards (red) induced a strong bias towards CCW responses, causing a leftwards shift of the function in addition to the shallower slope. The opposite bias arose with CW flankers (yellow). Both aspects of crowding are thus captured here: assimilative errors via the PSE, and the impairment in performance via threshold values.

**Figure 1.**
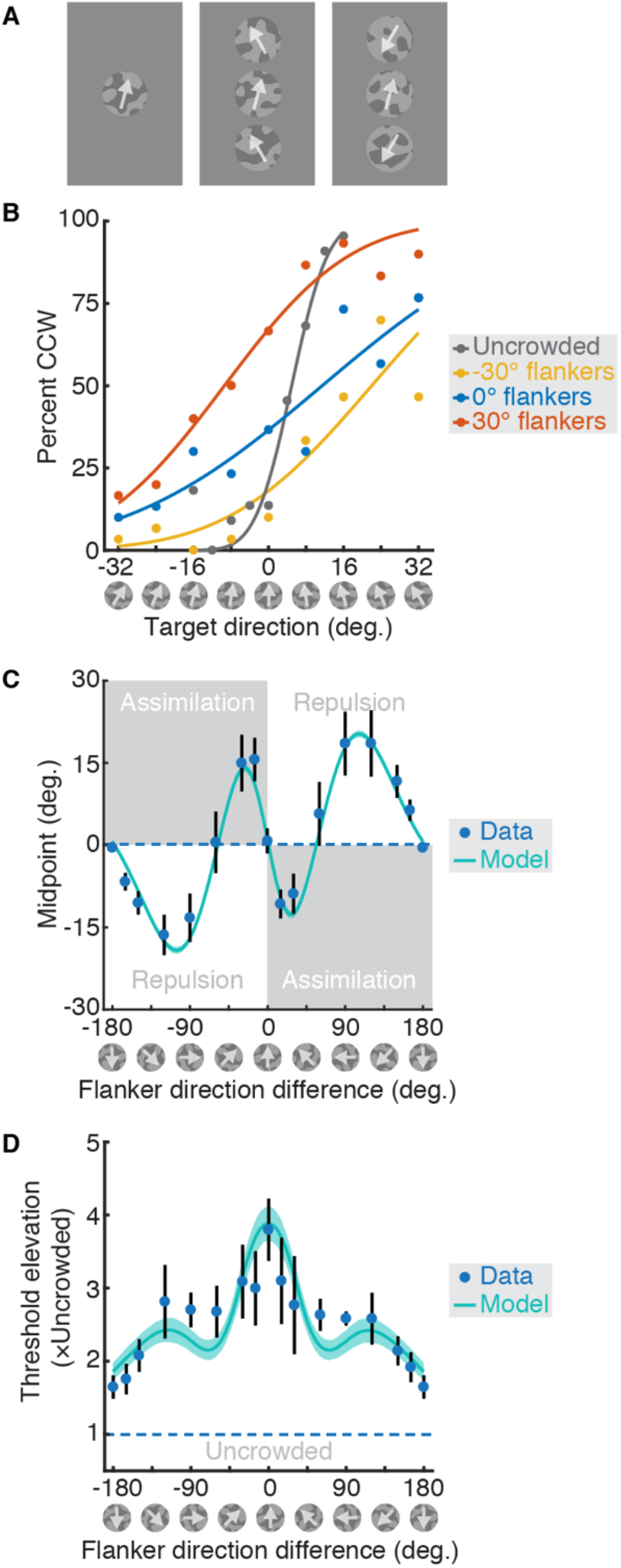
The effect of crowding on motion perception (Experiment 1). **A.** Left panel: An unflanked ‘cowhide’ stimulus. Middle: A crowded array with the target between flankers moving 30° CCW of upwards. Right: Crowded by flankers moving 150° CCW of upwards. **B.** Example data and psychometric functions for observer YL, with the proportion of CCW responses plotted as a function of target direction. Data are shown for an unflanked target (grey), and with flankers moving upwards (blue), −30° CW of upwards (yellow), and 30° CCW (red). **C.** Midpoint (PSE) values averaged over 6 observers (blue points with error bars ±1 SEM), plotted as a function of flanker direction. The mean output of a population crowding model is shown (green line) surrounded by the 95% range of values. **D.** Threshold elevation values for the same conditions, plotted as in panel C.

Psychometric functions were fit separately for each flanker condition and observer. Mean PSE values across observers are plotted as a function of the target-flanker difference in Figure 1C. On average, upwards moving flankers (0°) did not induce any bias, as with the example observer. Flankers moving slightly CW (e.g. −15°) induced a positive PSE shift, indicating an increase in CW responses. These assimilative errors were mirrored for small target-flanker differences in the CCW direction. Larger target-flanker differences (e.g. ±90°) induced a repulsive PSE shift, indicating that the perceived target direction was biased away from that of the flankers. Further increases gave a reduction in bias, with downwards flankers inducing no bias on average. Threshold elevation values (flanked thresholds divided by unflanked) are shown in Figure 1D, where a value of 1 indicates performance equivalent to unflanked thresholds (dashed line). The greatest threshold elevation occurred with upwards-moving flankers, with a decline in threshold elevation as flanker directions diverged. Downwards-moving flankers gave the least threshold elevation, though values remained above 1 for all observers. Altogether, crowding was strong with assimilative errors when target-flanker differences in motion were small, and reduced for large target-flanker differences with either repulsive errors or minimal biases.

We next examined the effect of crowding on judgements of hue in Experiment 2. Here observers identified whether the target was blue/turquoise or purple/pink (Figure 2A; Movie S2). When present, flankers differed from the reference hue by 1 of 12 hue angles in DKL colour space (26–28). Example data are shown in Figure 2B. Flankers with the same hue as the reference boundary (0°; blue points) did not induce any bias, though the slope is shallower than when unflanked (grey points). Flankers with a purple +15° hue angle (purple points) induced both a shallower slope in the psychometric function and a shift in the PSE, indicating assimilative errors, as did the blue −15° flankers (turquoise points).

**Figure 2.**
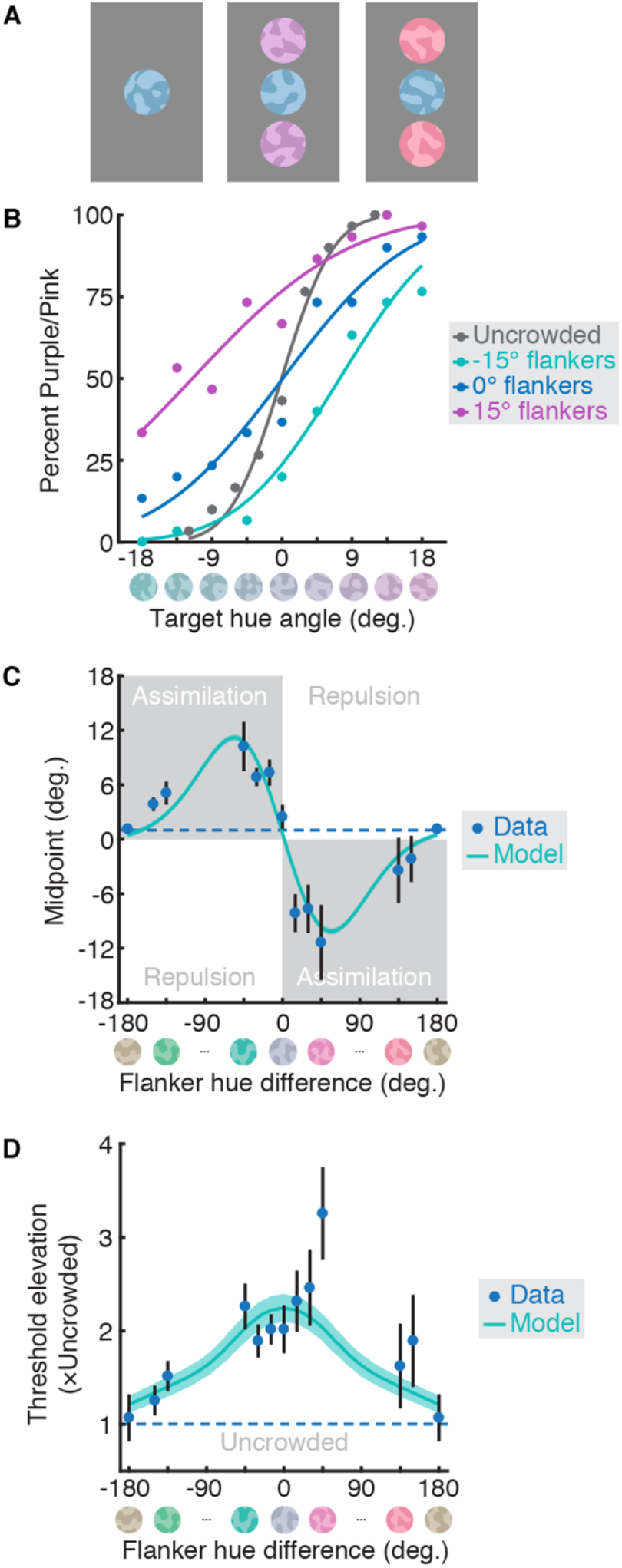
Crowding for colour perception (Experiment 2). **A.** Left: An unflanked stimulus. Middle: A flanked array with the target between flankers with +15° hues (purple). Right: A target with +135° flankers (pink/red). **B.** Example data and psychometric functions for observer AK, plotting the proportion of trials with a purple/pink response as a function of target hue (depicted on the x-axis). Data are shown for an unflanked target (grey), and flanked by stimuli with hues near the decision boundary (blue points), +15° CCW (purple), and −15° CW (turquoise). **C.** Midpoint (PSE) values averaged over 6 observers (blue points ±1 SEM), plotted as a function of flanker hue. The mean output of a population model of crowding is shown (green line) surrounded by the 95% range of values. **D.** Threshold elevation values for the same conditions, plotted as in panel C.

Figure 2C plots the mean PSE values for all flanker conditions. As with motion, flanker hues at the decision boundary (0°) induced no bias on average. Flankers with CW hue differences (blue-to-green in appearance) also induced positive shifts in PSE, indicating an increase in ‘blue’ responses. Assimilative errors were again mirrored for flankers with CCW hue angles, ranging from purple/pink through to red, while larger target-flanker differences gave little-to-no assimilative bias. Unlike motion, no errors of repulsion were observed. Mean threshold-elevation values are shown in Figure 2D. Although threshold elevation values are lower than for motion, the pattern of data is broadly similar, with the greatest threshold elevation for small target-flanker differences and a decrease in crowding strength with increasing difference. Flankers with the greatest differences (yellow/brown hues) did not elevate thresholds relative to unflanked performance.

Altogether, the crowding of both motion and colour is selective for target-flanker similarity – threshold elevation is high with small target-flanker differences and low with larger differences. In both cases, crowding also produced systematic errors that were predominantly assimilative for small target-flanker differences and declined with larger differences (though direction errors were repulsed at intermediate differences, which was not apparent for hue). More generally, the results of both experiments are broadly consistent with observations that biases follow the derivative of squared thresholds in a range of perceptual domains (29).

With this knowledge, we can now make predictions for paired judgements of motion and colour. Namely, when crowding is strong for one feature (with small target-flanker differences, e.g. in direction) and weak for the other (with larger differences, e.g. in hue), independent crowding processes allow assimilative errors to occur for the feature with strong crowding, without errors in the other. In contrast, a combined mechanism predicts that crowding must be all-or-none: if crowding is weak for one feature then it must be either reduced or persist for both.

Experiment 3 was designed to distinguish these alternatives: observers made conjoint judgements of the direction (CW/CCW of upwards) and hue (blue/pink) of the target cowhide for isolated targets and in three crowding-strength conditions. In the first, crowding was strong for both features, with small target-flanker differences in direction and hue (Movie S3). In the second, crowding was weak for direction (large direction difference) and strong for hue (small hue difference; Movie S4). The third involved strong crowding for direction (small differences) and weak crowding for hue (large differences; Movie S5). Each crowding-strength condition had 4 combinations of target/flanker elements with respect to the decision boundary for each feature dimension: either both motion and colour matched (e.g. CW moving target and flankers, all blue in hue), motion differed (e.g. a CW target with CCW flankers, all purple), colour differed (e.g. a purple target with blue flankers, all moving CW), or both differed. The crucial condition is when ‘both differ’: the all-or-none combined mechanism predicts errors in either both features or neither, while the independent mechanism allows a reduction in crowding in one feature without affecting the other.

With an unflanked target, observers correctly identified its direction in 87.71 ±3.29% (mean ±SEM) of trials, and its hue in 93.96 ±1.76% of trials. Figure 3A shows mean responses for the first crowding-strength condition, with strong crowding for both features. When target and flankers were matched in both feature dimensions (red point), performance was high in both cases. Here, even if crowding occurred, the assimilative effect of the flankers would pull responses toward the correct direction/hue. In the *motion differs* condition, observers were largely correct on the hue and incorrect for direction. This again is predicted by assimilative errors for direction, with either no effect on hue or assimilative crowding towards the correct hue. The converse occurred for the *colour differs* condition, with a predominance of colour errors. Finally, in the *both differ* condition, the strong assimilation for direction and hue induced errors for both features.

**Figure 3.**
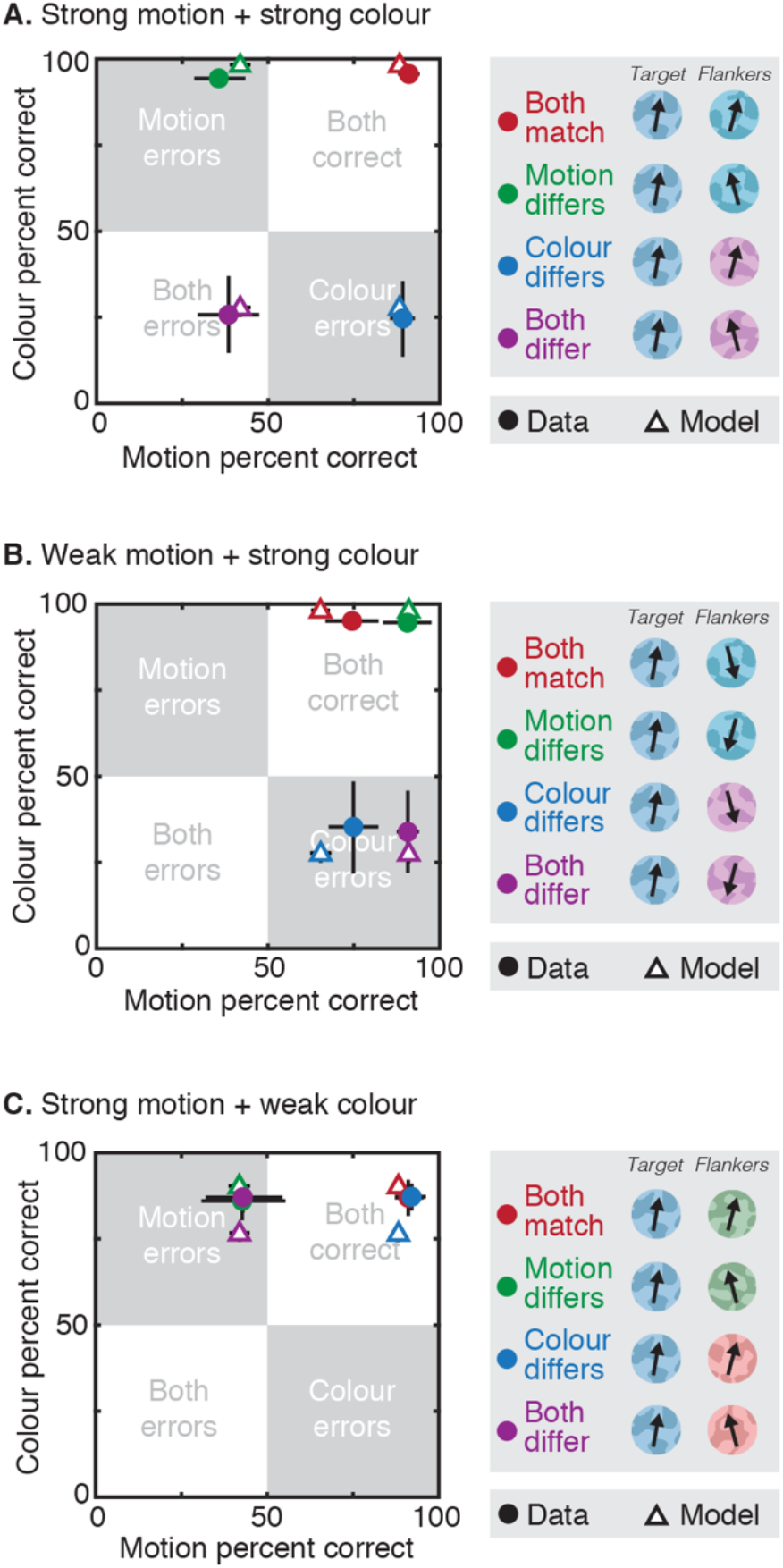
Results from the conjoint crowding of motion and colour (Experiment 3). Data (circles) is plotted as the mean proportion correct (n=6) for the target direction (x-axis) and hue (y-axis) ±1 SEM. The mean output of the best-fitting independent crowding model (triangles) with separate weights for motion and colour is also shown ±1 SEM. Quadrants are demarcated to show the predominant error type (e.g. ‘motion errors’). In each crowding-strength condition (separate panels) there were 4 target-flanker match conditions: where the 2AFC sign was matched for both features, where the motion differed, colour differed, or both differed, as shown by the legend. **A.** Strong motion + strong colour crowding condition. **B.** Weak motion + strong colour crowding. **C.** Strong motion + weak colour condition.

Figure 3B shows results from the weak motion + strong colour crowding condition. As before, in the *both match* condition, responses were correct on both features. In the *motion differs* condition, the large direction difference gave a reduction in crowding, with predominantly correct responses for direction, and likewise for hue given the matched target-flanker colours. For *colour differs*, the small hue difference continued to induce assimilative errors, while the similar target-flanker directions gave either assimilative errors or correct target recognition. Crucially, in the *both differ* condition, responses were correct for direction (as in the *motion differs* condition) but errors remained for hue, shifting responses into the ‘colour errors’ quadrant. Overall, the reduction in motion errors causes data for all conditions to align along the x-axis, while the separation along the y-axis for colour is retained. In other words, crowding was weak for motion and strong for colour in the same stimulus.

The converse pattern can be seen in the strong motion + weak colour condition (Figure 3C). Responses were again close to ceiling in the *both match* condition. In the *motion differs* condition, the small target-flanker direction difference again induced a high rate of assimilative motion errors and low rate of colour errors. Here in the *colour differs* condition, the large colour difference reduced crowding for hue judgements, while the matched target-flanker signs for direction led to correct responses for both features. Finally, the *both differ* condition again revealed a dissociation – large differences in target-flanker hue coupled with a small difference in direction produced errors in direction responses despite correct responses for hue. The reduction in colour crowding thus collapses data along the y-axis, while the separation for motion errors on the x-axis is retained. Here too, crowded errors can occur for one feature and not the other.

These errors follow the prediction of independent crowding processes for motion and colour, and are inconsistent with the predictions of a combined mechanism, whereby errors should have clustered in either the ‘both correct’ or ‘both errors’ quadrants. Accordingly, although errors in the *both differ* condition appear correlated on a trial-by-trial basis when crowding is strong for both features (SI Appendix; Figure S1A), this correlation breaks down when crowding is reduced for either feature (Figures S1B and S1C). We have further replicated these results with an increase in crowding strength – with additional flankers, we find stronger modulation in the crowding of motion and colour (Figure S2), but the pattern of independent errors for conjoint judgements of the two features remains (Figure S3). Finally, we also report that these dissociations in crowding are not confined to motion and colour – conjoint judgements of luminance-contrast polarity and direction show that errors can be low for contrast polarity and yet remain high for the direction of the same stimulus (Figure S4).

### Models

To better understand the mechanisms underlying these errors, and for the quantitative comparison of combined vs. independent mechanisms, we developed a set of computational models. Given the systematicity of crowded errors in these experiments, the most plausible models are those based on averaging or substitution (7, 11). A more general approach has been shown to produce both averaging and substitution errors by combining population responses to target/flanker elements (30). To simulate motion crowding in Experiment 1, we therefore developed a model population of direction detectors, with responses to target and flanker directions combined according to a weighting field. Where prior studies have used weighting fields that decreased with target-flanker distance (30), here the weights altered crowding strength as a function of target-flanker dissimilarity. To simulate the observed repulsion errors, we incorporated inhibitory interactions between target and flanker population responses, similar to models of the tilt illusion (31, 32). Further details and best-fitting parameters are given in the SI Appendix.

The best-fitting simulations of the crowded biases for motion in Experiment 1 are shown in Figure 1C (green line). The model follows the increase in assimilative bias with small target-flanker direction differences (driven by summation of the target and flanker population responses), as well as the rise and fall of repulsion with larger differences (driven by inhibition of the target; SI Appendix Figure S5). Similarly, threshold elevation values (Figure 1D) show the greatest elevation for small direction differences, with a decline on either side.

A similar population model was developed for colour crowding in Experiment 2 (SI Figure S5). Given the lack of repulsion for colour, inhibitory model parameters were set to zero. Figure 2C plots simulated biases (green line), which again capture the strong assimilative errors with small target-flanker hue differences and their decrease at larger differences. Threshold elevation values are similarly well described (Figure 2D), with a strong impairment for small target-flanker differences that progressively declines. Population-coding models can thus capture the errors observed for colour as well as for motion.

We next used these population models to simulate the conjoint motion and colour judgements of Experiment 3. Given the independent pattern of errors observed, we focus here on the operation of an independent crowding model where responses to target/flanker elements were combined via separate weighting fields for direction and hue (see SI Appendix). The independence of these weights meant that the strength of crowding for one feature did not affect the other. Figure 3A shows the best-fitting simulations for this model in the strong motion + strong colour condition, which follow the pattern of data well because the probability of crowding is high for both features. The model performs similarly well in the weak motion + strong colour condition (Figure 3B) because the separate weights allow crowding to be decreased for motion but not colour, leaving a predominance of colour errors in the *both differ* condition. Similarly, in the *both differ* condition with strong motion + weak colour crowding (Figure 3C), errors decreased for colour but remained strong for motion. Overall, the model follows the observed pattern of errors closely.

We have also developed a range of ‘combined’ models that use the same weight to crowd motion and colour on each trial. As outlined in the SI Appendix, these models consistently fail to replicate the pattern of errors found in Experiment 3 (Figures S6 and S7). Variations of the combined mechanism do little to improve performance – models that use the minimum or maximum probability for crowding in both features, and those with a single weighting field all produce worse fits than the independent model. Regardless of the precise mechanism, the crowding of motion and colour is best explained by independent processes.

## Discussion

Our perception of motion and colour is disrupted by crowding. Here we show that these effects are dissociable, indicating that they derive from independent processes. In Experiment 3, observers made judgements of both features while we manipulated the strength of crowding separately for each, using values from Experiments 1 and 2. When crowding was weak for motion (via large target-flanker direction differences) and strong for colour (via small differences), errors were reduced for motion but remained high for colour. Similarly, a reduction in colour crowding did not reduce errors for judgements of the target direction. A population-coding model of crowding reproduced this double dissociation by pooling target and flanker signals with independent weights for motion and colour. Models where crowding operated as a combined all-or-none process (with matched crowding strength for both features) failed to replicate these results.

Dissociations were also evident in the crowded errors for motion and colour measured in Experiments 1 and 2. Firstly, the overall magnitude of biases and threshold elevation was lower for colour than for motion. This difference decreased with additional flankers (SI Figure S2), further suggesting that crowding increases with flanker number at different rates for the two features. Secondly, intermediate target-flanker differences in motion caused a repulsion in perceived target direction, while equivalent colour differences simply reduced the rate of assimilative errors. Our population models reproduced these patterns via inhibitory interactions for motion, which were absent for colour. This does not mean, of course, that contextual modulations for colour are never repulsive. Although similar contextual effects tend towards assimilation in the periphery (33), repulsion in the perceived hue of targets does occur in foveal vision (34). A progression from foveal repulsion to peripheral assimilation also occurs for orientation (9). Given that motion repulsion occurs in both foveal and peripheral vision (35), it may be that the progression from repulsion to assimilation is more rapid across eccentricity for colour than motion. In other words, these distinct patterns of crowded errors offer further support for independent processes, though they may reflect variations in a common principle.

Although these dissociations for motion and colour crowding are consistent with the separation between these features in the visual system (17, 18), our findings differ from prior studies using other feature pairs. We attribute this to the degree of separation between these features in the visual system. For instance, the mixed pattern of independent and combined errors with spatial frequency, colour, and orientation (23) may have arisen because colour is dissociable from orientation and spatial frequency (as suggested recently; 36), while orientation and spatial frequency are more closely linked. The combined pattern of errors found for orientation and position crowding (22) could similarly reflect the interdependence of these features (37). That is, features that are closely related in the visual system may show linked performance, while more distinct feature pairs give dissociable effects. Comparable patterns are evident in other visual processes – for instance, colour and orientation show independent decay rates in visual working memory, unlike more closely linked spatial dimensions (38). A strong feature association could similarly explain the release in crowding for spatial-form judgements by differences in colour or contrast polarity (16). In these cases, however, the spatial forms are typically defined by the differential features (i.e. the spatial distribution of colour/polarity gives both the object surface and its boundaries; 39), making the colour or polarity signals informative regarding the feature being judged. Dissociations may only become evident when features can be judged independently, as in the current study.

Importantly however, a single dissociation between features is sufficient to reject an object-selective mechanism. Our results rule out this mechanism with at least two dissociations: colour and motion (Figure 3), and contrast polarity and motion (SI Figure S4). These results are similarly inconsistent with higher-level theories of crowding. Gestalt approaches (21) argue that crowding occurs when the target is ‘grouped’ with the flankers, e.g. by forming a pattern with the flankers (40), and that it is reduced when the flankers form patterns that exclude the target (41). The top-down nature of grouping suggests that it should apply to the collection of features within the target as a whole, making it an all-or-none process that is inconsistent with the dissociations found here. Our findings are equally unlikely to be accounted for by attentional theories (19, 20) since the high-level nature of attentional selection predicts that crowding should operate at the level of objects or locations, rather than being divisible for specific features within a localised target. Of course, attention and grouping could certainly modulate the strength of crowding – our findings simply suggest that these processes are not central to crowding.

Our population-coding model of these effects is similar to prior approaches in crowding and related contextual modulations (30–32). Here we show their generalizability to the domains of motion and colour. In fact, the dissociable nature of crowding lends itself to this approach – distinct populations with independent weighting fields for these features require fewer assumptions than a combined mechanism (SI Figure S6). Population coding may also explain the above distinction between combined crowding errors with some feature pairs and independent errors with others – the separation between these features in a multi-dimensional space (driven perhaps by their cortical distance; 9, 42) could determine the nature of these target-flanker interactions. Of course, it is also possible that ‘texturization’ models (43–45) could reproduce many of these effects, though distinct spatial and temporal texture processes would be needed to reproduce the dissociations for motion and colour.

The dissociation between motion and colour crowding further suggests that they may rely on distinct neural substrates. The many neural correlates of crowding reported from V1 through to V4 (46–49) may in fact reflect this distributed nature. In the most minimal sense, crowding in the ventral stream (44) may differ from the dorsal stream processes (17) likely involved in the crowding of motion. Crowding effects for other dissociable feature pairs may then be similarly distributed. It follows that crowding may be more profitably viewed as a general property of the visual system, similar to distributed processes like adaptation that affect a range of visual features (50). It is also possible, however, that dissociations could arise within a single cortical region through the operation of distinct neural subpopulations (as argued for feature-binding processes; 51).

At first glance, the distributed basis of these crowding effects bears some similarity to multi-level theories of crowding (4). However, these theories are based on an apparent uniqueness in the crowding of faces (52, 53), an effect that disappears once task difficulty is equated for upright and inverted faces (54). Although we do observe some differences in the crowding of motion and colour (e.g. with repulsion for motion vs. pure assimilation for colour), the broad selectivity of crowding was nonetheless highly similar in Experiments 1 and 2. Namely, small target-flanker differences gave strong assimilative errors and high threshold elevation for both features, while large differences gave a reduction in threshold elevation. In other words, wherever crowding occurs, it follows similar principles.

One complication with this distributed view of crowding is the common size of interference zones observed across a range of visual features (Bouma’s law; 2, 3, 12). Although differences may yet emerge for the specific comparison of motion and colour, this common spatial region may again be consistent with our effects deriving from distinct neural subpopulations with varying featural selectivity but common spatial properties. Alternatively, the proximity of target and flanker signals on the cortical surface (9, 42) may determine their potential for interaction, while the specific features present determine the nature of these interactions.

Altogether, we demonstrate that crowding independently disrupts motion and colour, whilst nonetheless operating via common principles (seen in the implementation of our population models). This dissociation excludes the possibility that crowding operates as a singular mechanism and suggests that at least some aspects of vision are disrupted by clutter in a feature-specific manner.

## Materials and methods

### Observers

6 observers (3 male, including the authors) completed all 3 experiments. All had normal or corrected-to-normal acuity, and normal colour vision as assessed by the Ishihara test (55). Informed consent was given, with procedures approved by the Experimental Psychology ethics committee at University College London.

### Apparatus

Experiments were programmed in MATLAB (Mathworks, Inc.) on an Apple Mac Pro using the PsychToolbox (56, 57). Stimuli were presented on a 21” Mitsubishi Diamond Plus CRT monitor with a resolution of 1400×1050 pixels and 75Hz refresh rate. The monitor was calibrated using a Minolta photometer and linearised in software to give a mean and maximum luminance of 50 and 100 cd/m^2^, respectively, and a white point near the standard CIE Standard Illuminant D65. Maximum luminance values for red, green, and blue were 28.3, 69.5, and 8.1 cd/m^2^, respectively. Observers viewed stimuli binocularly from 50cm distance, with head movements minimised using a head and chin rest. Responses were given by keypad, with auditory feedback provided only during practice sessions.

### Stimuli and Procedures

In all experiments, target and flanker stimuli were ‘cowhide’ elements (24, 25), created by band-pass filtering white noise with a spatial frequency cut-off of 1.5cyc/deg, and rounding the luminance to give two values (light and dark). Each element was presented within a circular aperture with 2° diameter. The visible contours in these elements enabled the percept of motion with minimal ambiguity given their orientation variance (i.e. avoiding the aperture problem; 24), whilst also allowing alteration of the surface hue.

Observers were required to maintain fixation on a two-dimensional Gaussian blob with a standard deviation of 4’. The target was presented 15° above fixation, either in isolation or with one flanker above and one below. The centre-to-centre separation of target and flankers was 2.25°, corresponding to 0.15× the eccentricity (well within standard interference zones; 2, 3). Stimuli were presented for 500ms, followed by a mask for 250ms (a patch of 1/f noise in a circular window of diameter 4.8° when unflanked and 8.5° when flanked, plus a cosine edge). The mask was followed by a mean-grey screen with the fixation point, at which time observers responded.

In Experiment 1, cowhide stimuli were grey-scale elements with a Weber contrast of ±0.75 against the mean-grey background. Patches were generated as a long strip of texture that moved behind the aperture with a displacement of 5.8’ per frame every second monitor frame (to allow greater resolution of directional displacements with larger, less frequent steps). This gave an effective stimulus refresh rate of 37.5Hz and a speed of 3.6deg/sec.

When unflanked, the target moved in 1 of 9 equally spaced directions between ±16° around upwards and ±32° when flanked (given the greater difficulty). Observers indicated whether the target moved counterclockwise (CCW) or clockwise (CW) of upwards. When present, flankers moved together in 1 of 16 directions relative to upwards: 0, ±15, ±30, ±60, ±90, ±120, ±150, ±165, or 180°. Each block had 10 repeat trials per target direction, giving 90 trials for unflanked blocks and 180 for flanked conditions, where opposing flanker directions (e.g. ±15°) were interleaved within a single block to ensure a balanced likelihood of CW and CCW responses. 0° and 180° conditions were also interleaved for consistency. Each block was repeated 3 times, with all blocks randomly interleaved, to give 4590 total trials per observer, plus practice, completed in 3-4 sessions of 1 hour each.

In Experiment 2, cowhides were static and presented with a range of hues. Colours were determined using the DKL colour space (26–28) with a luminance contrast of ±0.3 for light and dark regions and a colour contrast/saturation of 0.2. Variations were applied solely to the hue angle. The reference hue angle was determined individually, given variation in the categorical boundaries for colour between observers (28). We did so by presenting the test range of hues (from blue/turquoise to pink/purple) and asking observers to indicate the neutral midpoint. This gave a reference hue of 262.5° for four observers, 262.0° for JG, and 264.0° for CS. When unflanked, the target was presented with 1 of 9 equally spaced hues ±12° from the base hue, and from ±18° when flanked. Observers judged whether the target appeared blue/turquoise (CW in DKL space) or purple/pink (CCW). When present, flankers had 1 of 12 hue angles relative to the base: 0, ±15, ±30, ±45, ±135, ±150, and 180°, tested in blocks that contained opposing angles as above. This gave 90 trials per unflanked block and 180 when flanked, giving 3510 total trials per observer, plus practice, completed in 3 sessions.

In Experiment 3, cowhides varied in both direction and hue. For each observer, we selected values from the first two experiments that gave near-ceiling performance levels when unflanked but that were clearly impaired by crowding in the strongest crowding conditions. For direction, this gave values of ±5° (YL), ±6° (CS and JG), ±7° (DO), ±10° (AK), and ±16° (MP), and for hue ±3° (CS and YL), ±4° (DO), ±5° (JG), ±7° (AK), and ±10° (MP). Observers indicated the direction and hue of the target as a 4AFC response – either Blue/CCW, Blue/CW, Pink/CCW, or Pink/CW. Targets were either presented in isolation or with flankers selected for each feature to give either ‘strong’ or ‘weak’ crowding (as above). Strong flanker directions were ±10° (DO), ±15° (AK, CS, JG, and YL), or ±30° (MP), with weak values of ±165° for 5 observers or ±175° (AK). Strong flanker hues were ±15° (AK and YL) or ±30° for the remainder, with ‘weak’ values of ±150° for 5 observers or ±165° for JG.

In addition to the unflanked condition, the above combinations of target-flanker elements gave 3 crowding-strength conditions: strong motion + strong colour crowding (small target-flanker differences for each), weak motion + strong colour crowding (large motion, small colour differences), and strong motion + weak colour crowding (small motion, large colour differences). For flanked conditions, there were 16 combinations of direction and hue values in the target and flanker elements (2 target directions ×2 flanker directions ×2 target hues ×2 flanker hues). We grouped these conditions into 4 combinations of target/flanker elements in terms of their agreement in the 2AFC decision space for each feature. In the *both match* conditions, both motion and colour were matched in target and flanker elements. When *motion differed*, the sign of the target direction differed from that of the flankers (e.g. a CW target with CCW flankers) but their hues matched. Conversely, when *colour differed* the hue of the target differed from the flankers, while directions were matched. Finally, target and flanker elements could *both differ* in their direction and hue values. Note that these distinctions relate to the decision boundary, ignoring precise values of direction/hue (e.g. −15° and −165° flankers have the same sign as a −8° target). The four crowding-strength conditions were tested in separate blocks, with each combination of target/flanker elements repeated 10 times per block to give 40 trials when unflanked and 160 trials for flanked conditions. Each block was repeated 6 times, randomly interleaved, with 3120 total trials per observer (plus practice), completed in 3 sessions.

### Analyses

In Experiments 1 and 2, psychometric functions were fit to data as a cumulative Gaussian function with 3 free parameters: midpoint/PSE (at 50%), slope, and lapse rate. Functions were fit separately for each flanker condition and observer. Shifts in the midpoint were taken as changes in appearance (i.e. assimilation vs. repulsion errors). Thresholds were taken as the difference in direction/hue required to shift performance from the midpoint to 75% CCW responses, with threshold elevation obtained by dividing flanked by unflanked thresholds. Data in Experiment 3 were combined from the 16 target-flanker combinations into 4 target-flanker match conditions, and analysed as the percent correct in each feature dimension, treating each as a 2AFC judgement.

### Models

Data in Experiments 1 and 2 were fit with a population-coding model based on that by Harrison & Bex (30). The motion-crowding model of Experiment 1 had 9 free parameters, with 5 free parameters for the colour model in Experiment 2 (since the lack of repulsion allowed inhibitory components to be removed), as described in the SI Appendix (Figure S5). Table S1 shows the best-fitting parameters, with final outputs in Figures 1 and 2. In Experiment 3, the independent model for motion and colour crowding involved population responses to target and flanker elements that were combined via separate weighting fields for each feature. The majority of parameters were carried forward from Experiments 1 and 2, leaving 3 free parameters (Table S2). Outputs of the best-fitting model are in Figure 3. A series of combined models were also developed, which were identical to the independent model, save for the use of common weights for both features. Tables S3 and S4 show the best-fitting parameters, with outputs in Figures S6 and S7.

### Data availability statement

SI datasets are available in proportion CCW format for each observer in Experiments 1 and 2, and proportion correct for each observer in Experiment 3. MATLAB code for psychometric functions and stimulus generation is available at http://github.com/eccentricvision.

## Acknowledgements

Funded by a UK Medical Research Council Career Development Award (MR/K024817/1). Thanks to Alexandra Kalpadakis-Smith, Samuel Solomon, and Christoph Witzel for helpful comments. Parts of this work were presented to the Vision Sciences Society (58).

## Supporting Information Appendix

### Trial-by-trial correlations in the crowding of motion and colour

We demonstrate in Experiment 3 that crowded errors for motion and colour are dissociable. Central to this experiment was the *both differ* condition, where target and flanker elements had opposite signs relative to the decision boundary (e.g. a clockwise-moving blue flanker amongst counterclockwise-moving purple flankers). In this condition when crowding was strong for both motion and colour, errors were high for both features. In contrast, when crowding was strong for motion and weak for colour, *both differ* responses were predominantly incorrect for motion and correct for colour. Conversely, with weak motion and strong colour crowding, responses were correct for motion and incorrect for colour. These dissociations are consistent with independent crowding processes for motion and colour.

Another way to approach this issue is to examine trial-by-trial variations. If crowding were driven by a combined all-or-none crowding mechanism, responses on individual trials should be either incorrect for both features (due to crowding) or correct on both (when crowding is released). In contrast, independent crowding mechanisms allow trial-by-trial dissociations in the same way as observed for error rates across the whole experiment. We therefore sought to test these predictions by examining the correlation between responses for motion and colour across trials.

Responses on each trial were initially encoded as binary variables (incorrect vs. correct). We first determined the overall proportion of each response type across the experiment by converting the response on each trial to one of four outcomes – responses were either incorrect on both motion and colour, correct on both, correct on motion but incorrect on colour, or vice versa. Proportions were calculated for each observer in each condition, collapsed across both target and flanker sign (e.g. whether the target was moving CW or CCW). Here we report the outcome of these analyses for the *both differ* condition only, given that this was the condition required to separate performance of the independent and combined models.

This classification provides a 2×2 table of outcomes. The mean proportion of each response across observers in the strong motion + strong colour condition is shown in Figure S1A. Similar to the percent-correct values shown in Figure 3, responses were most frequently incorrect for both features in a given trial, with colour errors forming the second-most frequent error type, closely followed by errors in both features. In this case then, the high proportion of errors in both features appears consistent with a trial-by-trial correlation. To quantify this further, we computed *phi* coefficients (used to quantify correlations for binary variables) using the binary response outcomes on each *both differ* trial, separately for each observer. These correlations were significant for 5/6 observers (AK: ϕ=.217, *p*<.001, CS: ϕ=.378, *p*<.0001, DO: ϕ=.228, *p*<.0001, JG: ϕ=.187, *p*=.004, MP: ϕ=.458, *p*<.0001, YL: ϕ=.024, *p*=.705). In other words, an error for motion was likely to coincide with an error for colour. Of course, the correlated outcome in this condition derives from the high strength of crowding in both features (coupled with the tendency for correct responses on both features in around 20% of trials). Both models can therefore explain this outcome.

Compare now the results for the *both differ* condition with weak motion + strong colour crowding (Figure S1B). A combined model predicts that observers should either be correct for both features, or incorrect on both. In contrast, the most likely outcome was a correct motion and incorrect colour response, followed by correct responses for both features at half the rate, and negligible proportions in the remaining cells. The correlation between these responses on a trial-by-trial basis was not significant for any of the observers (AK: ϕ=.092, *p*=.154, CS: ϕ=−.125, *p*=0.053, DO: ϕ=.042, *p*=.514, JG: ϕ=0, p=1, MP: ϕ=.019, *p*=.767, YL: ϕ=.025, *p*=.700). In other words, the response shifted to a predominance of errors in colour rather than motion for all of our observers, consistent with the predictions of an independent model.

**Figure S1.**
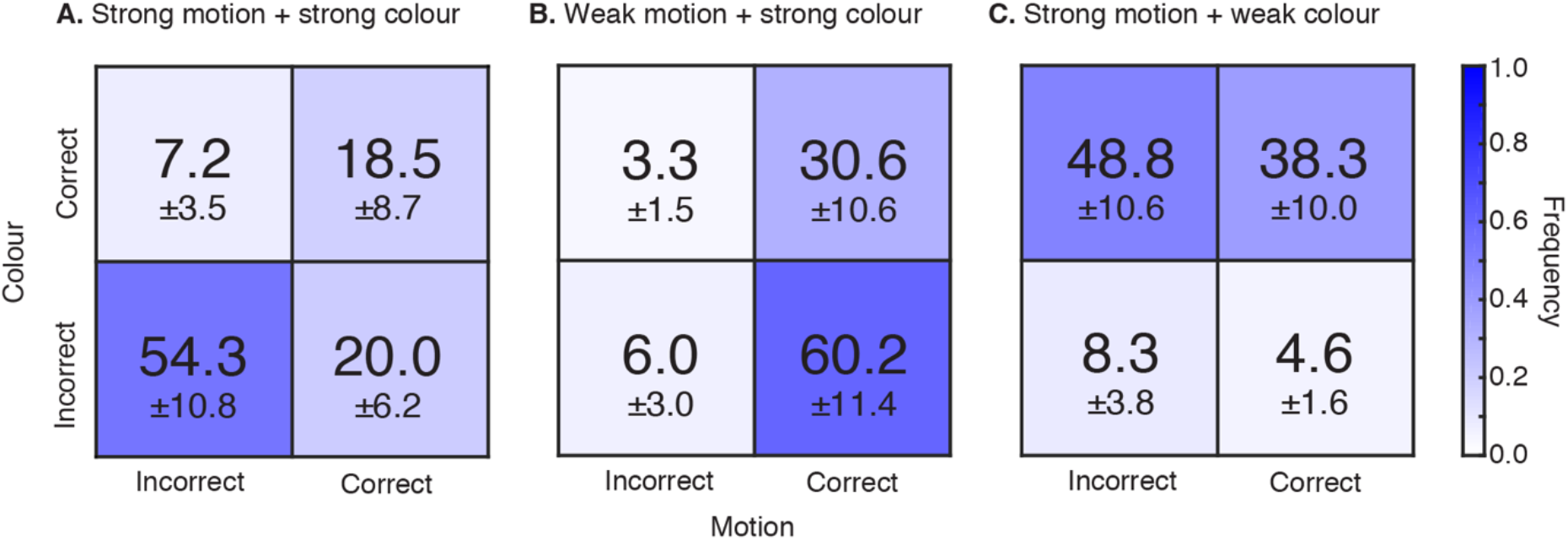
The frequency of response errors across the trials of Experiment 3. Data from the *both differ* condition is shown for the strong motion + strong colour crowding condition (panel A), the weak motion + strong colour crowding condition (panel B) and the strong motion + weak colour crowding condition (panel C). Data is reported as the mean proportion of trials in which each of the four error types occurred, as indicated both by the colour of each cell (see colour map) and numerically ±1 SEM.

Finally, results for the *both differ* condition with strong motion + weak colour crowding are shown in Figure S1C. Here the most likely outcome was an incorrect motion and correct colour response, followed by correct responses for both features, and negligible proportions in the remaining cells. The correlation between these responses was not significant for 5/6 observers, with a negligible correlation for the remainder (AK: ϕ=.034, *p*=.593, CS: ϕ=.044, *p*=.495, DO: ϕ=.120, *p*=.064, JG: ϕ=.080, *p*=.216, MP: ϕ=−.080, *p*=.214, YL: ϕ=.058, *p*=.028). This predominance of errors in motion rather than colour is again consistent with the predictions of an independent model.

Altogether, when crowding was strong in both features, the high proportion of errors on each gave a significant trial-by-trial correlation. In contrast, responses were clearly uncorrelated in the latter two crowding-strength conditions given that a reduction in crowding for one feature did not affect the rate of errors in the other. These findings are again consistent with independent crowding processes for motion and colour.

### Experiments S1-S3: Crowding for motion and colour with increased flanker numbers

Experiments 1-3 reported in the main text used two flankers to induce crowding, positioned along the radial axis with respect to fixation. We selected this configuration because radial flankers have the strongest influence on crowding (59), and because the effect of radial vs. tangential flankers on target appearance can vary depending on their contour alignment with the target (60–62). It is possible however that other configurations could increase the strength of crowding, and in turn that combined effects of crowding on motion and colour judgements may become more apparent with this increased strength. Because crowded performance impairments have been shown to increase in conjunction with the number of flankers (63, 64, though cf. 65), we repeated our experiments with four flankers.

Another potential complication in Experiments 1-3 is with our use of a post-stimulus mask – in each experiment, a patch of 1/f noise was presented immediately after stimulus offset in order to minimise visual persistence (66). It is possible however that these masks interfered with the stimulus. In particular, if the masks were to interfere more strongly with one feature than the other (e.g. colour; 67), this may have promoted independent effects in our data. We therefore removed the post-stimulus mask for these experiments.

To assess whether these manipulations altered crowding strength, whilst also ensuring that large target-flanker differences continued to give a sufficient reduction in crowding, we examined the crowding of motion and colour separately with abridged versions of Experiments 1 and 2, respectively (Experiments S1 and S2). We then used these parameters to measure conjoint judgements of motion and colour, as in Experiment 3 (Experiment S3). Six observers were tested (4 female), including one of the authors (JG), one who participated in the prior experiments (AK) and four new naïve observers. A seventh observer was excluded given threshold values for direction that were outside the measurable range.

Experiment S1 measured motion crowding under these circumstances. As above, the number of flankers was increased to four, positioned above, below, left and right of the target when present (Figure S2A). The post-stimulus mask was also removed, with participants responding immediately after stimulus offset. Because our aim was to find one target-flanker difference with strong crowding and another with weak crowding (as parameters for the conjoint Experiment S3), we also reduced the number of flanked conditions from 16 to 4. Based on the results of Experiment 1, observers were thus presented with a small target-flanker difference (2 conditions, ±15° from upwards) expected to produce strong crowding and a large difference (±175°) expected to give reduced crowding. Note that the latter difference was increased from Experiment 1. Observers were presented with these 4 flanked conditions in 2 blocks, plus a third block for the unflanked condition, with 3 repeats of each interleaved randomly. Target directions ranged from ±20° in 11 steps in the unflanked condition and ±40° in 11 steps for flanked conditions. This range was increased from Experiment 1 given observer reports of the difficulty in these conditions. Observers began with practice blocks with 2 repeats per target direction, repeated until performance stabilised. They then began the final testing phase with 10 repeats per direction. Testing took 1-2 hours in hour-long sessions, including practice. Remaining stimulus parameters and procedures were identical to those of Experiment 1 in the main text.

As before, data were first analysed as percent counterclockwise responses, with psychometric functions fit to the data to obtain bias and threshold estimates. Given our aim to determine the strength of crowding in each condition, here we further converted bias values to ‘assimilation scores’ by reversing the sign of biases for clockwise flanker conditions. This made positive and negative values indicative of assimilation and repulsion, respectively. Values within each flanker difference condition (±15 and ±175) were averaged for each observer. Mean assimilation scores across observers are shown in Figure S2B. Here the small target-flanker difference led to strong assimilative bias, with a mean of 21.08°, reducing to a small degree of repulsion at the larger separation, with a mean of −2.73°. For comparison, the equivalent conditions in Experiment 1 (±15° and ±165°) gave assimilation scores of 13.18° and 6.52°. The manipulations in Experiment S1 were therefore successful both in increasing the strength of crowding with small target-flanker differences, as well as increasing the reduction in assimilative errors with larger differences.

**Figure S2.**
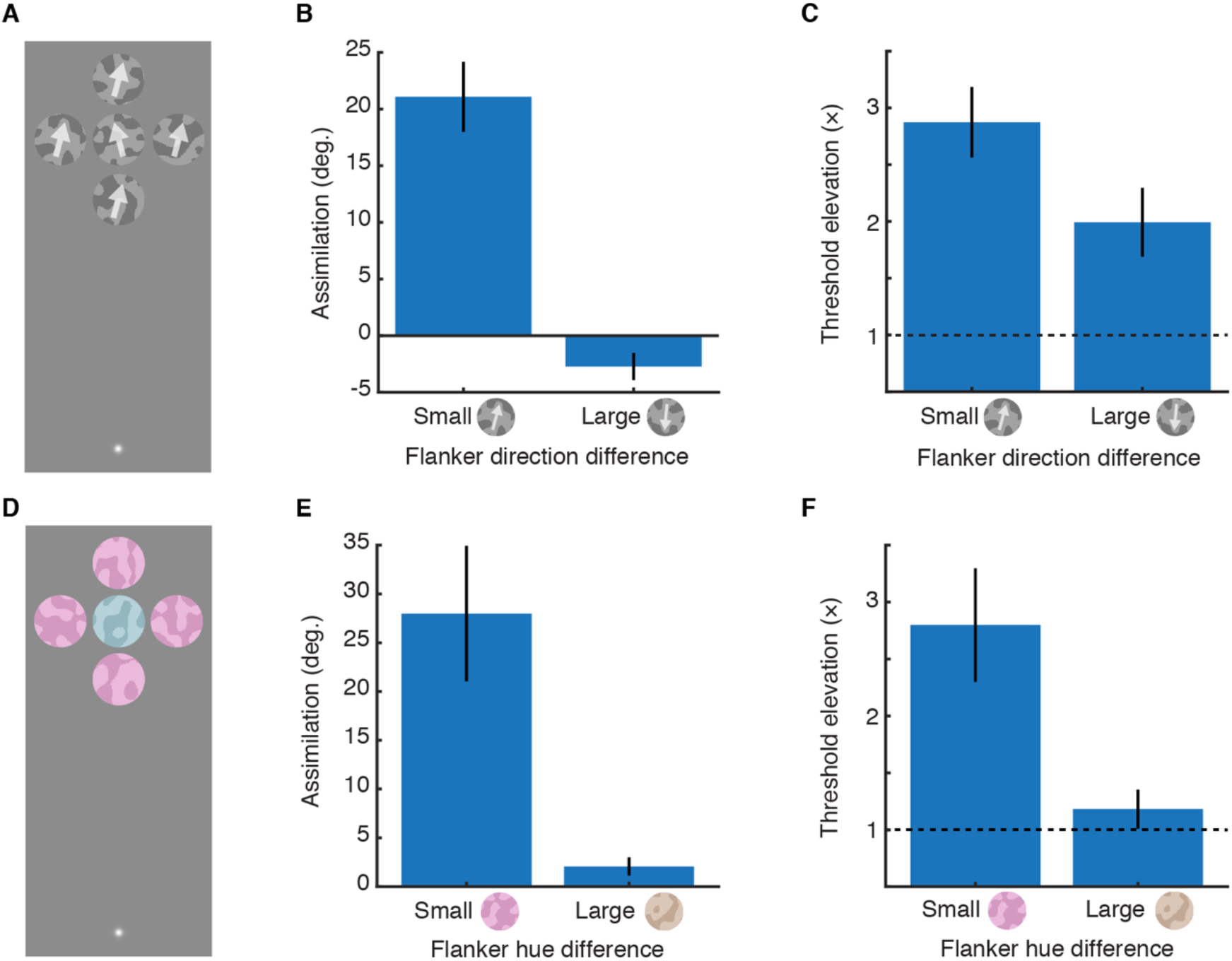
Results for Experiments S1 and S2 with 4 flankers. **A.** Example stimuli from Experiment S1 with a small directional offset in the flankers. **B.** Assimilation scores for motion crowding in Experiment S1 with small (±15°) and large (±175°) direction differences in the flankers. Positive values indicate assimilation and negative values repulsion. **C.** Threshold elevation scores for Experiment S1, plotted as multiples of unflanked thresholds. **D.** Example stimuli from Experiment S2 with a small hue difference in the flankers. **E.** Assimilation scores for colour crowding in Experiment S2 with small (either ±15° or ±30°) or large (±175°) hue angle differences in the flankers. **F.** Threshold elevation values for Experiment S2.

Threshold values were similarly averaged across clockwise and counterclockwise conditions for each of the two flanker difference conditions, before being divided by unflanked thresholds to give threshold elevation values, plotted in Figure S2C. Here the small target-flanker difference gave thresholds 2.87× higher than unflanked levels, while thresholds with the large difference were 1.99× higher. Equivalent values from Experiment 1 were 3.05 and 1.83. Here then there is little change in threshold elevation from two to four flankers, though the slight increase (for both target-flanker separations) is consistent with observer reports that the target was occasionally difficult to see under these conditions. This is likely consistent with the previously observed increase in the impairment of detection thresholds as flanker number increases (63). Nonetheless, in conjunction with the bias values reported above, crowding effects were more strongly modulated here than in Experiment 1.

Experiment S2 was then conducted to examine similar values for the crowding of colour. As above, we conducted this experiment with four flankers (Figure S2D), the removal of the post-stimulus mask, and a reduction in the number of target-flanker difference conditions. During practice blocks, two observers (AK and MH) showed only minimal crowding effects with hue differences of ±30° and were therefore tested with ±15° for the small target-flanker difference condition (consistent with values used in Experiment 3). The large target-flanker hue difference was ±175° for all observers (giving hues that were rusty orange or yellow/brown in appearance). Target hue differences ranged from ±15° in 11 steps in the unflanked condition and ±25° in 11 steps for flanked conditions, again increased from the values of Experiment 2 given the higher difficulty with this configuration. For simplicity all observers were set to have the same decision boundary for hue (262.5°), unlike the variations in Experiment 2 (which had only minimal effect). The remaining stimulus parameters and procedures were identical to Experiment 2.

Data were analysed in the same way as Experiment S1, with mean assimilation scores shown in Figure S2E. On average, small hue differences in the flankers induced a strong assimilative bias of 27.98°, whereas large differences gave biases of only 2.06°. This brings colour biases in line with those seen above for motion, presenting a marked increase from the corresponding values in Experiment 2 (using the values of ±15/30° and ±150/165° selected for Experiment 3), which were 7.02° and 2.74°, respectively. Similarly, thresholds were elevated by 2.80× and 1.18× unflanked levels for the small and large differences, respectively (Figure S2F). This too is an increase on the values of 2.34 and 1.47 in Experiment 2. In other words, this configuration gave more bias and higher threshold elevation with small target-flanker differences, as well as a larger decrease in these values with large target-flanker differences.

Given this increase in crowding strength, we next used these parameters with conjoint judgements of motion and colour in Experiment S3. Observers were tested with the flanker values used in Experiments S1 and S2, again with 4 flankers and the removal of the post-stimulus mask. Target direction and hue values were selected as in Experiment 3 as values that were gave near-ceiling performance in the unflanked condition whilst also being clearly shifted by biases in the flanked conditions. This gave target directions of ±8° (for AS, MF, MH, and JG) and ±10° (for AM and HC) around upwards, and hue differences of ±5° (for MH), ±8° (for AS, MF, HC, and JG) and 10° (for AM) from 262.5°. Remaining parameters were identical to those of Experiment 3.

With an unflanked target, observers correctly identified its direction in 87.64 ± 1.45% (mean ± SEM) of trials, and its hue in 90.00 ± 2.59% of trials. Figure S3A shows mean responses for the flanked condition with small target-flanker differences in each feature (to give strong crowding for both). When the target and flankers were matched in sign for both features (red point, e.g. a purple CW target amongst purple CW flankers), performance was again high in both cases. This could be due either to a lack of crowding or the assimilative effect of the flankers pulling responses toward the correct direction/hue. In the *motion differs* condition, observers were largely correct on the hue and incorrect for direction. This again is predicted by assimilative errors for direction, with either no effect on hue or assimilative crowding towards the correct hue. The opposite occurred for the *colour differs* condition, leading to a predominance of colour errors. Finally, the *both differ* condition induced a high rate of errors in both features, indicative of strong assimilation for each.

Figure S3B shows results from the weak motion + strong colour crowding condition. As before, in the *both match* condition, responses were clearly correct on both features. In the *motion differs* condition, the large direction difference gave a reduction in crowding, with predominantly correct responses for direction, and likewise for hue given the matched target-flanker colours. For *colour differs*, the small hue difference continued to induce assimilative errors, while the flanker directions gave either assimilative errors or correct target recognition. Crucially, in the *both differ* condition, responses were correct for direction (as with *motion differs*) but errors remained for hue, shifting responses into the ‘colour errors’ quadrant. In other words, crowding was strong for colour and weak for motion in the same stimulus.

**Figure S3.**
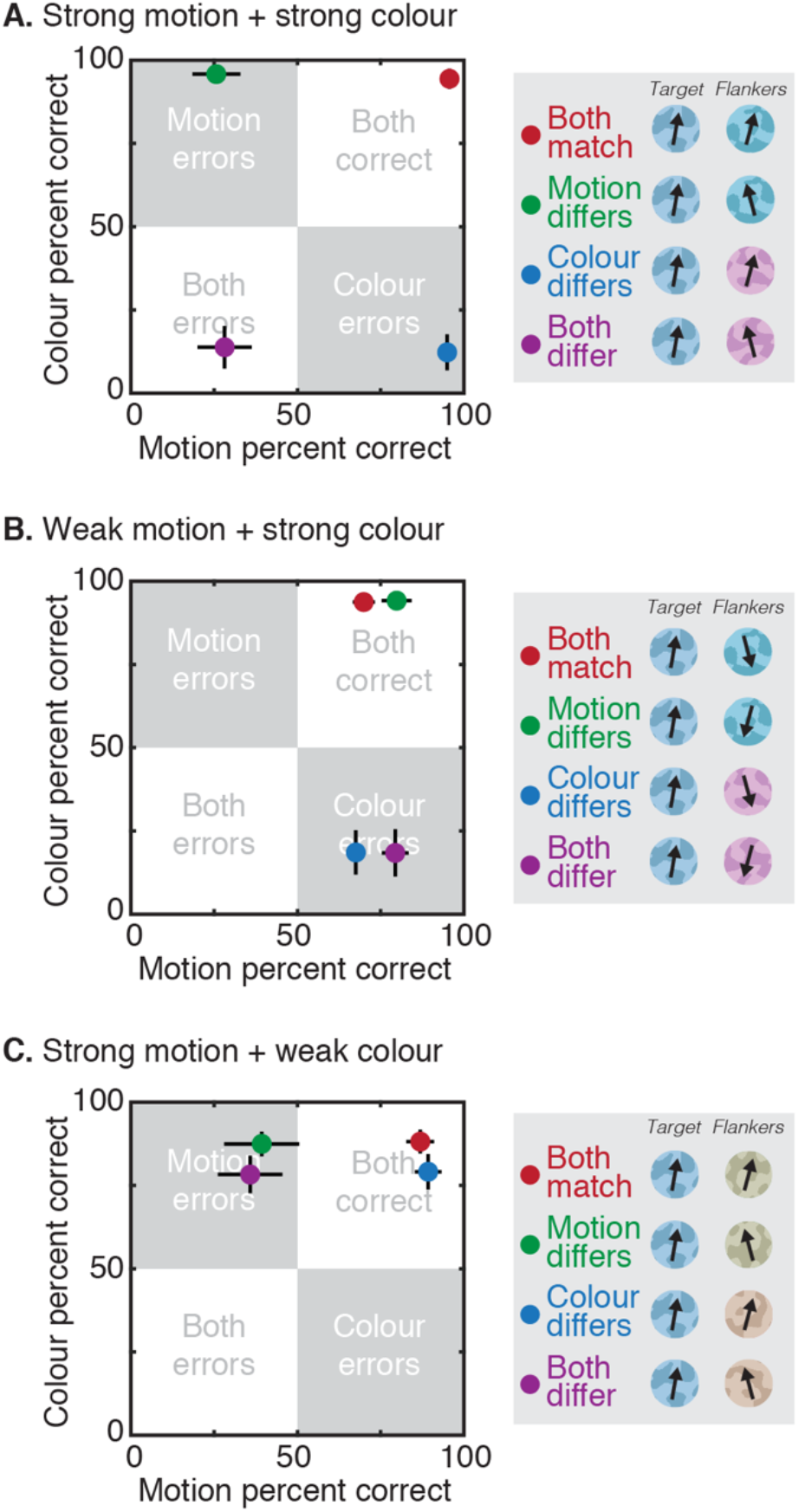
Results from the conjoint motion and colour judgements of Experiment S3 with 4 flankers. Data is plotted as the mean proportion correct for the target direction (x-axis) and hue (y-axis) ±1 SEM, as in Figure 3 of the main text. Indicative target-flanker values are shown via the legend. **A.** Results from the strong motion + strong colour crowding condition. **B.** Results from the weak motion + strong colour crowding condition. **C.** Results from the strong motion + weak colour condition.

The reverse pattern can be seen in the strong motion + weak colour condition (Figure S3C). Responses were again close to ceiling in the *both match* condition. In the *motion differs* condition, the small target-flanker direction difference induced a high rate of assimilative motion errors, with a low rate of hue errors. Here in the *colour differs* condition, the large colour difference reduced crowding for hue judgements, while the matched target-flanker signs for direction led to correct responses in both features. Finally, the *both differ* condition again revealed a dissociation – small target-flanker differences in direction coupled with large differences in hue produced consistent errors in direction despite correct responses for hue. Here too, crowding can occur for one feature and not the other.

These errors again follow the prediction of independent crowding processes for motion and colour, and replicate our earlier result with an increased number of flankers. An increase in the strength of crowding did not therefore change the independence of these errors. In fact, the difference between performance in the target-flanker match conditions is somewhat larger than that shown in Figure 3 (e.g. along the y-axis in the weak motion + strong colour crowding condition), given the greater assimilative effects for colour in particular. This replication also demonstrates that our use of a post-stimulus mask in Experiment 3 did not artificially produce an independent pattern of errors.

The differential effect of flanker number observed these experiments offers further support for the independence of motion and colour crowding. In particular, although the increase to four flankers gave a modest increase in biases and threshold elevation for motion in Experiment S1 relative to those in Experiment 1, the average rate of assimilative errors for colour more than tripled in Experiment S2 from the values in Experiment 2. Observers also noted that the judgements of motion in Experiment S1 were highly difficult, with an occasional tendency for the target to disappear, an effect that was not found in earlier experiments (corroborated by AK and JG who completed all experiments). This could be due to an increase in the suppressive effect of the flankers as their number increases, seen also with the increased repulsive effect in Experiment S1 with large direction differences. Suppression could also explain the lower increase in assimilative biases for motion relative to Experiment 1, compared with the strong increase in the purely assimilative biases for colour. Nonetheless, this divergence in the effect of flanker number for motion and colour offers further evidence that independent crowding processes affect these two features.

### Experiment S4: Crowding for motion and luminance-contrast polarity

Our results demonstrate that crowding is independent for judgements of direction and hue. It is possible, however, that this finding is somehow specific to these two feature dimensions. A common manipulation in studies seeking to reduce crowding is the use of target-flanker differences in luminance contrast polarity – a black target amongst white flankers (or vice versa) gives considerably less crowding than elements with uniform polarity (16, 68, 69). Here we sought to test the generality of our conclusions with conjoint judgements of direction and luminance contrast polarity.

Unlike hue, luminance contrast polarity is a binary property (light or dark), with linear variations in luminance contrast between these extremes. It is therefore not ideal to run our full paradigm with this feature – the conjoint judgements of Experiments 3 and S3 involve fine discriminations around a decision boundary with flankers that are either close to this boundary (small differences leading to strong crowding) or distant (leading to weak crowding). Here, variations in luminance contrast around the decision boundary would fall close to zero contrast (mean grey), affecting the visibility of these stimuli. This in turn would likely introduce errors in the motion judgements, given issues with target detectability, potentially making errors appear combined simply because the target is invisible. We can nonetheless run a subset of these conditions with the maximum values of luminance contrast (i.e. full white vs. full black) and examine the effect of reductions in crowding from contrast polarity on judgements of direction.

In Experiment S4 we therefore required observers to make conjoint judgements of direction and contrast polarity. As in prior experiments, a combined mechanism for crowding predicts that any reduction in crowding for contrast polarity must also reduce crowding for direction. In contrast, independent mechanisms allow for errors in direction to remain high even with a reduction in crowding for contrast-polarity judgements.

The design of this experiment was similar to that of Experiments 3 and S3. Here, cowhide elements varied in both direction and luminance contrast polarity. For this purpose, we rendered only half of the cowhide elements (unlike the combined light/dark regions in other experiments), with one half of the image rendered either white or black, and the remainder left as the mean grey of the background (Figures S4A and S4B). The same 6 observers who completed Experiments S1-S3 also took part in this experiment. Target values for direction were taken from those used in Experiment S3: ±8° around upwards for AS, MF, MH, and JG and ±10° for AM and HC. Luminance contrast polarity values were set at their maximum value of ±1, giving a Weber contrast of 100% against the mean grey background.

Observers made conjoint judgements of the direction (CW/CCW of upwards) and contrast polarity (black/white) of the target cowhide for unflanked targets and in one flanked condition (strong motion + weak contrast polarity). Following Experiment S3 above, the flanked condition included 4 flankers (above, below, left and right of the target). When flankers were present, their directions were similar (±15°) in order to induce strong crowding for motion. Contrast polarity values were set to their maximum contrast difference (±1) to give maximally strong crowding when target and flanker elements were matched in polarity and maximally weak crowding when polarity differed. Given the above issues with the dimensionality of contrast polarity, we did not test the converse arrangement with large directional differences. Unflanked and flanked conditions were tested in separate blocks. Each combination of target and flanker elements was repeated 10 times per block to give a total of 40 trials in the unflanked condition and 160 trials for the flanked condition. Each block was repeated 3 times, with all blocks interleaved randomly.

With these combinations of direction and contrast polarity, there were 4 possible combinations of target and flanker elements with respect to the decision boundary for each feature dimension: either both motion and contrast polarity matched (e.g. CW moving target and flankers, all black), motion differed (e.g. a CW target with CCW flankers, all white), contrast polarity differed (e.g. a white target with black flankers, all moving CW), or both differed. Given the known effects of differences in contrast polarity on crowding (16, 68, 69), judgements of contrast polarity should be correct in conditions where this property differs (‘polarity differs’ and ‘both differ’). They should also be correct when target and flanker elements share the same contrast polarity (‘both match’ and ‘motion differs’), either because there is no crowding or because responses are driven by assimilative errors. The crucial aspect of this experiment is the motion judgements, particularly in the ‘both differ’ condition. Here the all-or-none combined mechanism predicts errors in either both features or neither, while the independent mechanism allows a reduction in crowding in contrast polarity without any effect on motion (i.e. that motion errors should remain high).

**Figure S4.**
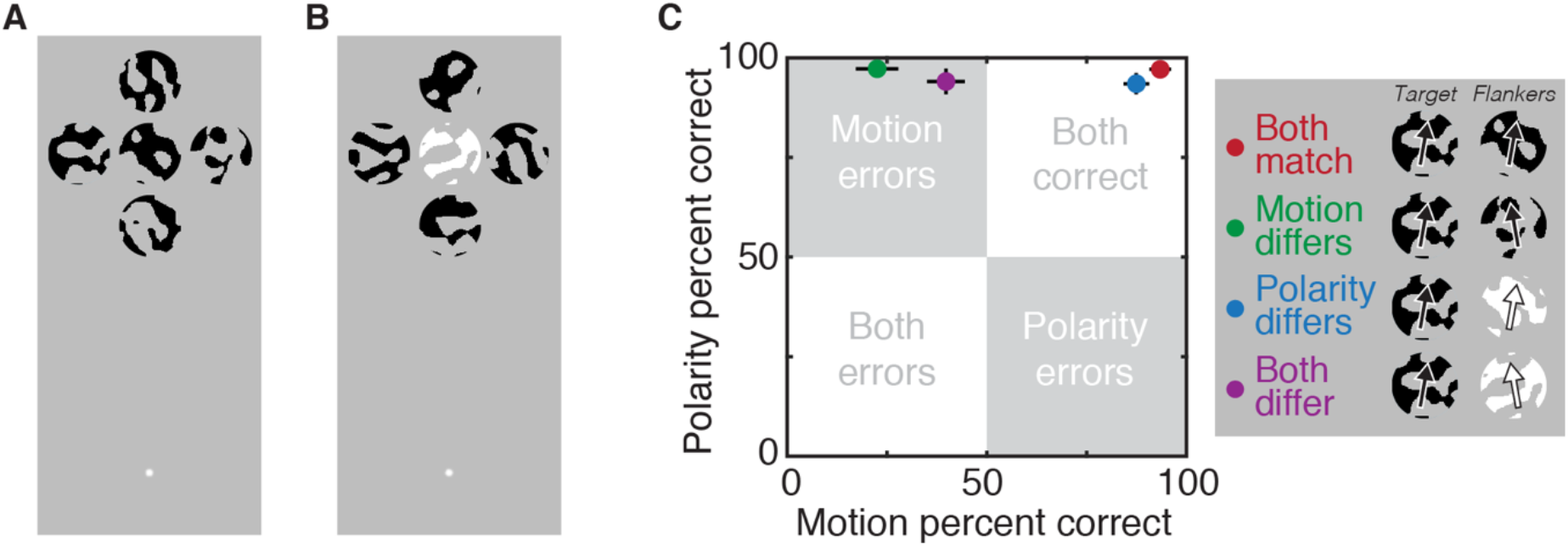
Stimuli and results for Experiment S4 examining judgements of motion and luminance contrast polarity. **A.** Example stimuli when all black. **B.** Example stimuli when contrast polarity differed between the target (here, white) and flankers (black). **C.** Results from the strong motion + weak contrast polarity condition. Data is plotted as the mean proportion correct for the target direction (x-axis) and contrast polarity (y-axis) ±1 SEM, as in Figure 3 of the main text, with example target-flanker values shown via the legend.

With an unflanked target, observers correctly identified its direction in 83.89 ± 3.12% (mean ± SEM) of trials, and its contrast polarity in 98.19 ± 0.82% of trials. Figure S4C shows mean responses for the flanked condition, where target-flanker differences were small in motion (to give strong crowding) and large in contrast polarity (to give a release when target and flanker elements differed). When target and flankers were matched in both features (red point), performance was high for both features. In the *motion differs* condition, the small target-flanker direction difference induced a high rate of assimilative motion errors and low rate of contrast polarity errors, again because target and flanker elements remained matched in polarity. Judgements of polarity continued to be accurate in the *polarity differs* condition, given the large contrast-polarity difference (replicating prior findings with polarity differences; 16, 68, 69), while the matched target-flanker signs for direction allowed correct responses in both features. Crucially, the *both differ* condition again revealed a dissociation – the large differences in target-flanker polarity coupled with a small difference in direction produced errors in direction responses despite correct responses for contrast polarity. In other words, here too crowded errors can occur in one feature and not the other.

Interestingly, there is a slight increase in the percent correct for motion in the *both differ* condition, relative to the error rate for the *motion differs* condition in this experiment (i.e. a separation along the x-axis). This difference was not present in Experiments 3 or S3. One possibility is that the opposite polarity elements did in fact induce some degree of combined errors, perhaps through a reduction in positional uncertainty associated with the target location (70). Alternatively, several observers noted that opposite-polarity targets would on occasion disappear, which may relate to our use of four flankers in this experiment, and the rise in issues related to detection as the number of flankers increases (63), as noted above. This may be exacerbated at the 15° eccentricity used herein (particularly in the upper visual field; 19, 71), as most studies that have examined the release from crowding with opposite polarity stimuli have utilised closer eccentricities of 5-10° (16, 68, 69). Regardless, the reduction in these errors still leaves a predominance of motion errors, in contrast to the clear performance levels (above 90% correct) for contrast polarity judgements – motion judgements never approach this level of performance.

The observed pattern of errors again follows the prediction of independent crowding processes for motion and luminance contrast polarity, extending our findings with direction and hue. This finding does however stand at odds with prior demonstrations that crowding is reduced for judgements of spatial form (like T orientation) when target and flanker elements differ in contrast polarity (16, 68, 69). As outlined in the main discussion, the linkage found in these prior studies could reflect the greater similarity between contrast polarity and spatial form than between polarity and motion. Indeed, contrast polarity and motion have been found not to interact at higher levels of the motion hierarchy (72). Features that are more closely related in the visual system, like orientation and position (22), may therefore show linked performance patterns, while more distinct feature pairings like direction and hue or direction and polarity allow these dissociable effects to emerge.

Note also that prior studies showing a combined release for polarity and form (16, 68, 69) typically utilise spatial letterforms (e.g. T elements) defined by the differential features themselves. That is, the colour/polarity signals define both the object surface and its boundary (39), making the spatial distribution of luminance-polarity signals informative regarding the feature being judged. This may then allow the differences in polarity to reduce crowding for spatial form simply because the form signals for the target are derived from the (uncrowded) output of the polarity channels. The same is true for studies showing a release in orientation crowding with colour differences (13, 16, 73). Dissociations may only become evident when features can be judged independently, as in the current study where these features apply only to the object surfaces (given that our circular object boundaries were always held constant).

### Population models for the crowding of motion and colour

As shown in the main text, data from Experiments 1 and 2 were fit with two population-coding models similar to prior models of the crowding of orientation (30, 74). This approach characterises crowding as the weighted combination of population responses to the target and flanker elements, which has previously been found to reproduce the systematic errors that arise (30), including both averaging (7, 11) and substitution (75) errors. Here we sought to extend this modelling approach to the domains of motion and colour perception. To replicate the results of Experiment 1, we simulated a population of 361 direction-selective neurons, each with a wrapped Gaussian profile of responses to direction, similar to those found in cortical areas V1 (76) and MT/V5 (77), characterised as:

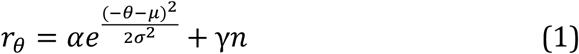

where r_θ_ is the response of the detector for a given value of θ, the direction ranging from ±180 around upwards. The value α sets the height of the tuning function (set to 1), μ is the direction of peak response, σ represents the standard deviation of the Gaussian (the first free parameter), plus Gaussian noise *n* with a magnitude of γ (the second free parameter). Responses outside the range ±180 were wrapped by either subtracting or adding 360 to the direction and summing the responses. Each of these detectors had a distinct preferred direction at one of the integer directions from ±180°. Flanker population responses had the same form (including the same standard deviation), with a separate free parameter for the γ value, representing a late noise parameter.

Given the presence of repulsion in our data (similar to the pattern of direction repulsion more broadly; 35, 78), we followed models of the tilt illusion (31, 32, 79, 80), and the physiology of MT/V5 neurons (81), by adding inhibitory surrounds to the population response. This is also similar to weighted averaging models of crowding that simulate repulsive errors using negative weights (60). Here we incorporated inhibitory interactions via a second Gaussian distribution (as in equation 1), with a peak of 0.3 for the population responses to the target and 1.0 for flankers (to be modulated by flanker weights, below), with the standard deviation of this distribution as the fourth free parameter. This distribution was then subtracted from the excitatory Gaussian response described above. The peak of 0.3 was selected given physiological estimates that place the strength of inhibition at around 30-40% that of excitation in early visual cortex (82). Values of 0 and 0.5 for the target population were also simulated, which did not alter the pattern of results in a qualitative fashion (though parameters varied to accommodate this inhibition of the flankers).

Population responses were determined for both the target and flanker directions separately. These responses were combined according to weights, as in prior models (11, 22, 30, 74, 83). Variations in these weights have previously been used to reproduce the decrease in crowding with increasing target-flanker distance (9, 30, 74) via ‘weighting fields’. Target-flanker distance was fixed in our study, though we utilise this weighting-field concept to allow the decrease in crowding with increasing target-flanker dissimilarity (9, 10, 13, 16). In doing so, we follow suggestions that both of these properties may in fact manipulate the cortical distance between target and flanker representations (9, 42, 84). Because our population used both positive and negative components, two weighting fields were applied separately (similar to prior work; 9). Positive weights varied from 0-1 and were determined using a Gaussian distribution (as in equation 1, though without noise), set as a function of the target-flanker *difference* in direction (rather than absolute direction above), centred on a difference of 0. The peak and standard deviation were each set by free parameters. Negative weights also varied from 0-1 and were set by a bimodal Gaussian distribution of the form:

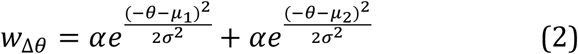

where *w* gives the flanker weight for a given difference in direction (Δθ), while the peak α and standard deviation σ were matched for each Gaussian. The peak was set as a free parameter, as well as the difference between the peak locations (μ_2_−μ_1_), with the overall distribution centred on zero. The standard deviation was the same value used in the positive weighting field. The shape of this function allowed for the peak in inhibitory interactions at large target-flanker differences where repulsion effects were dominant over assimilation (see Figure 1C). Overall however, both components varied with the direction difference between target and flanker elements to modulate the strength of these crowding effects. The best-fitting weighting fields are plotted in Figure S5A, which plot the change in flanker weights as a function of the target-flanker difference in direction. The corresponding target weight was always 1 minus the flanker weight for both positive and negative components.

For a given trial of the simulated experiment, flanker weight values were first selected according to the difference between target and flanker directions. These values were then used as multipliers for the positive and negative components of the population response to the flanker direction. Altogether then, flanked responses *C* were determined as function of the direction θ, with the form:

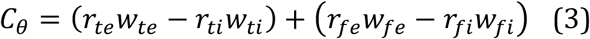

where *r*_*te*_ represents the excitatory Gaussian population response to the target (following equation 1), *r_ti_* is the inhibitory response, and *r*_*fe*_ and *r*_*fi*_ the excitatory and inhibitory flanker responses, respectively. Weight values are denoted as *w*_*fe*_ for the excitatory flanker values and W_fi_ as the inhibitory weight. For the target *w*_*te*_ is 1−*w*_*fe*_ and *w*_*ti*_ is 1−*w*_*fi*_.

This combination of responses and weighting values for the population response to flankers is plotted in Figure S5B for a continuous range of target-flanker differences (flanker values tested in Experiment 1 are shown with black points). Responses are plotted as a function of the preferred direction of each detector in the population on the x-axis against the target-flanker difference on the y-axis. The population response to a single flanker direction can be seen by taking a horizontal slice across the plot, with red areas indicating a predominance of positive flanker responses and blue areas indicating a predominance of inhibition. Note that small target-flanker differences near to the 0° decision boundary tend to produce predominantly positive flanker responses, whereas larger target-flanker differences tend towards inhibition.

Example population distributions (averaged across 1024 trials) for a target moving 8° counter-clockwise from upwards and flankers moving 90° counter-clockwise are shown in Figure S5C. Distributions of target and flanker responses (red and blue lines, respectively) have had their respective weights applied. The combined sum is shown in yellow. Veridical values of the target and flankers are shown as red and blue triangles. Notice that the target and flanker directions are both counter-clockwise, yet the peak response for the combined distribution lies on the clockwise side at −9° (yellow triangle) to produce an error of repulsion. This occurs due to the greater inhibition from the flankers on the counter-clockwise side of the population. In contrast, target-flanker combinations where the population response to the flankers was predominantly positive tended to induce assimilation effects by shifting the peak of the combined response to intermediate values between the target and flanker directions (see demonstration for colour below).

**Figure S5.**
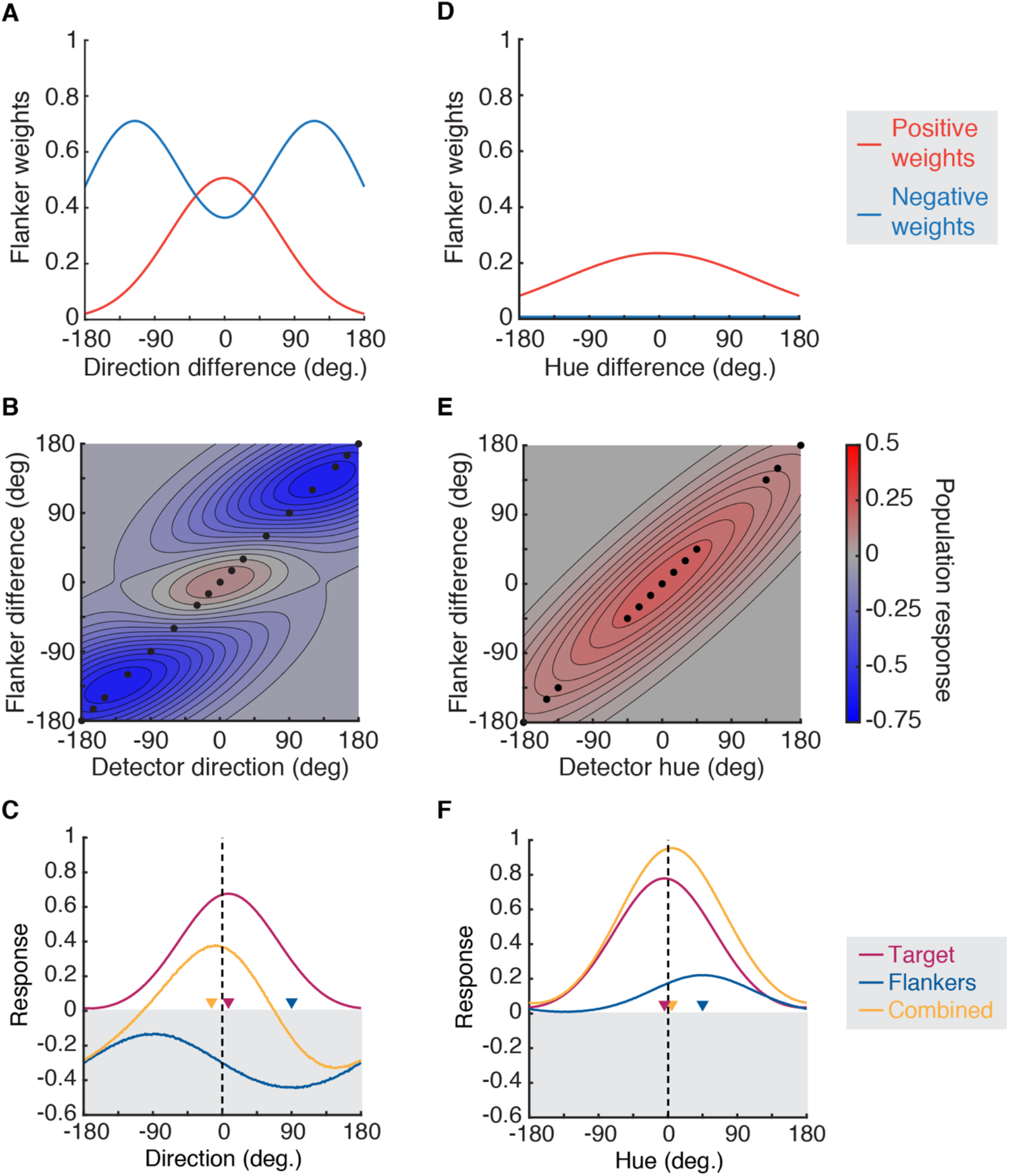
Model characteristics for the population-pooling models of Experiments 1 (left column) and 2 (right). **A.** Weighting fields for direction, plotted as a function of the target-flanker direction difference, separately for the positive (red) and negative (blue) weights. **B.** The combination of flanker weights and the population response to the flanker direction, plotted as a function of the preferred direction of each detector on the x-axis and the target-flanker direction difference on the y-axis. Flanker values tested in Experiment 1 are shown as black points. Red values indicate a predominance of positive population responses, while blue areas indicate inhibition. **C.** Example population responses to the target (red), flankers (blue), and combined response (yellow) for a target moving 8° counter-clockwise from upwards and flankers moving 90° counter-clockwise, plotted as a function of the preferred direction of each detector on the x-axis. The veridical values of the target and flanker directions are shown as red and blue triangles, with the peak response of the combined distribution shown as a yellow triangle. This combination gives a repulsion error. **D.** Weighting fields for hue in Experiment 2, plotted with conventions in panel A. **E.** The combination of flanker weights and the population response to hue, plotted as in panel B. **F.** Example population responses for a target with a hue −4.5° clockwise from the decision boundary (blue in appearance) and flankers 45° counter-clockwise (pink in appearance), which gives an error of assimilation. Plotting conventions as in panel C.

The perceived target direction was derived from the peak response of this combined population response distribution (equation 3) on each of the simulated trials, with the sign of this response used to determine the 2AFC (CW/CCW of vertical) response. Target and flanker direction conditions were identical to those of Experiment 1, with 1024 trials per condition in the simulation. As with the behavioural responses, the percent CCW was then computed for each target direction in each flanker direction condition, with psychometric functions fit to determine midpoint and threshold values. The squared difference between these midpoint and threshold values was then taken from the mean behavioural data of Experiment 1 to give an error term. The best-fitting parameters (for the above 8 free parameters) were determined first using a coarse grid search through the parameter space to find the least squared error, which was then used as the starting point for a fine fitting procedure using the *fminsearch* function in MATLAB. Best-fitting parameters from this procedure are shown in Table S1, with the output of the model plotted against the data in Figure 1 of the main text.

A similar model was developed to account for the pattern of responses to the colour task of Experiment 2. Because repulsion was not present in the data obtained from this study, all inhibitory components of this model (including target and flanker responses, as well as weighting fields) were set to zero, leaving 5 free parameters. The structure of the model was otherwise identical to that for motion, with a population of 361 hue-selective neurons with a Gaussian profile of responses to the hue angle in DKL colour space (26–28). Each detector had a preferred hue angle at one of the integer values in the space with a Gaussian tuning function on either side, consistent with suggestions from both physiological (85) and psychophysical results (86), and is used here for ease of comparison across features.

The best-fitting weighting field for colour is shown in Figure S5D. Notice that the absence of inhibitory weighting fields meant that the sole effect of crowding in the domain of colour was to induce assimilative errors. This can be seen with the combination of the weighting field and flanker population responses across a range of flanker hue angles plotted in Figure S5E. As before, these values were produced by combining the flanker response to a continuous range of target-flanker differences with the weighting field values for those same target-flanker differences (with the flanker values tested in Experiment 2 presented as black points).

**Table S1.**
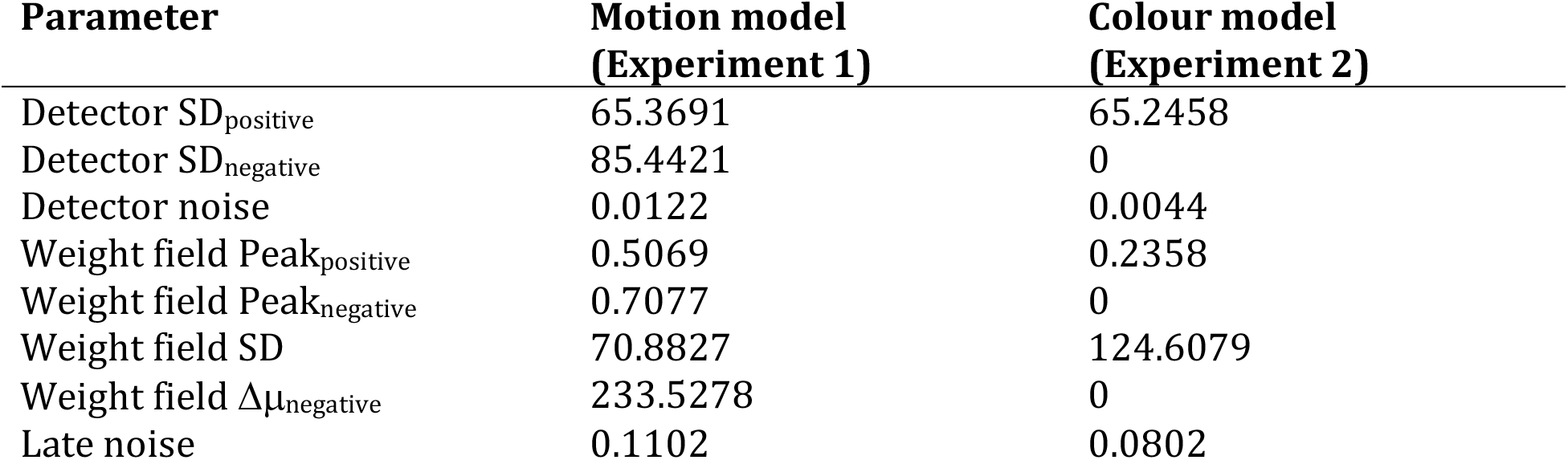
Best-fitting free parameter values for the population-based crowding model for motion (Experiment 1) and colour (Experiment 2).

As with motion, the combined population response was generated by summing the responses to target and flanker directions. Example population distributions for a target with a hue angle −4.5° clockwise from the blue/purple decision boundary, and flankers with a 30° counter-clockwise hue are shown in Figure S5F. When combined (yellow line) the assimilative nature of crowding can be seen – here the peak response for the combined distribution lies on the counter-clockwise side at 2° (yellow triangle). Best-fitting parameters (for the 5 free parameters) were determined using the same procedure as for motion, and are presented in Table S1, with the output of this model plotted against the data in Figure 2 of the main text.

Figures 1 and 2 show that the best-fitting output of these models provides a good characterisation of the pattern of errors observed for motion and colour crowding, and particularly in the case of motion crowding. We note that prior models have used a similar difference-of-Gaussian approach to model errors of repulsion in the domain of orientation, with mixed results (31, 32, 79, 80). The improved performance of the model in the present study likely derives from our addition of a weighting field (30), which allows a smooth transition from predominantly assimilative errors at small target-flanker differences through to strongly repulsive errors with larger differences (for motion, at least). Because the weighting field modulates the noise introduced by flankers (the ‘late noise’ parameter), this approach can also replicate the rise and fall of threshold elevation seen in Figures 1C and 2C. Although the use of these weighting fields will likely need adjustments to account for the many complexities of crowding (87), here we show their generalizability to the domains of motion and colour. In this way we also reproduce the general coupling between bias and threshold observed in a range of perceptual contexts (29).

### Independent population models for the crowding of motion and colour

The results of Experiment 3 followed our predictions for independent crowding processes for motion and colour. Here we quantify these processes with a population coding approach. The independent model consisted of population responses to motion and colour, generated for both target and flanker elements and combined according to separate weighting fields for both features. These separate weighting fields allowed for crowding to occur for one feature (with small target-flanker differences, e.g. in colour) and not in the other (with larger target-flanker differences, e.g. in motion). The majority of model properties were carried forward from Experiments 1 and 2, including the standard deviation of detector tuning functions, as well as the peak height and standard deviation of the positive weighting field, as in Table S1. Inhibitory parameters were included for the motion population. This left 3 free parameters: early noise for direction, early noise for hue, and the combined late noise parameter for both features. Note that the latter noise parameter applied to the flanker population responses (which was then combined with the target population response). Since this was modified by the weights for each feature, we used a single parameter for both features to reduce the number of free parameters and for greater ease of comparison with the combined models.

Because the precise direction and hue values varied between participants in this experiment (see values in the main text), we used the modal value for each as the input for the model. This gave a target direction difference of ±8° and a hue angle difference of ±5°. For direction, flankers differed by ±15° and ±165° for the strong and weak motion crowding conditions. Flanker hue angles were ±30° and ±150°. Each trial simulated the population response to target and flanker values for both motion and colour.

For the independent model, separate weighting fields were used to convert the target-flanker differences in direction and hue into flanker weights. These were identical to those of the first two experiments. Weights were applied to target and flanker population responses to generate a combined population response for each feature. Each peak response was then used to determine whether responses would be CW/CCW for motion and for hue, with percent correct determined across trials.

The 3 free parameters were fit by determining the least-squared error between the percent correct scores for motion and colour in each of the four crowding-strength conditions (unflanked, strong motion + strong colour, weak motion + strong colour, and strong motion + weak colour). Performance was simulated with 1024 trials per point. As before, a coarse grid search was conducted through the parameter space prior to a fine fitting procedure. Final parameters are shown in Table S2, with the output of this model plotted in Figure 3 of the main text.

**Table S2.** Best-fitting free parameter values for the independent crowding model and an alternative version of this model without inhibitory parameters, both used to simulate the data of Experiment 3.

| Parameter | Independent model | Independent model<br>(no inhibition) |
| --- | --- | --- |
| Direction noise | 0.0126 | 0.0353 |
| Colour noise | 0.0057 | 0.0038 |
| Late noise | 0.0408 | 0.0713 |

For the strong motion + strong colour condition, shown in Figure 3A, the independent model follows the pattern of data well because the probability of crowding (with two weighting fields) is high for both features, producing assimilative errors. In the weak motion + strong colour condition (Figure 3B), the model successfully captures the pattern of performance because the separate weighting fields for the two features allow crowding to be independently decreased in motion, leaving colour errors in the *both differ* condition. Conversely, the model again reproduces performance in the strong motion + weak colour condition (Figure 3C) because crowding can be reduced for colour and remain strong for motion, leading to a high rate of motion errors when *both differ*. It is therefore plausible that human performance could rely on independent crowding mechanisms of this nature.

In the following section, we outline a range of combined crowding models in an attempt to more quantitatively rule out the combined mechanism. Several of these approaches remove the inhibitory aspects of the model for simpler comparison across the feature dimensions. In order to more directly compare these models with the independent crowding model, we also simulated the above independent model with inhibitory parameters set to zero (both in population responses and the corresponding weighting fields). Best-fitting parameters are shown in Table S2. Mean squared error values for 1000 simulations of the Independent model without inhibition was 0.1086, slightly worse than the value of 0.0752 obtained for the above model with inhibition. The removal of inhibition thus impaired performance of this model to some extent, though both versions of the independent model vastly outperformed any of the combined mechanisms tested below (see Figure S7).

### Combined population models for the crowding of motion and colour

The results of Experiment 3 and associated simulations demonstrate that crowding is most likely to be subserved by independent processes for motion and colour. In order to more comprehensively rule out the possibility that a combined mechanism could perform similarly well in these experiments, we simulated a range of models with this combined all-or-none mechanism. These models were similar in operation to the independent models described above, save for the use of common weights for the two features. Here we show that these models all fail to fully account for the observed dissociations in motion and colour crowding.

Some assumptions must be made in developing a combined mechanism for two features. If we begin with a model that is otherwise identical to the independent approach described above, then motion and colour estimates are derived from each population of detectors for both target and flanker elements. Responses to the target and flankers must then be combined with weights. Consider first a model that retains the two best-fitting weighting fields derived from Experiments 1 and 2, but which takes the minimum value obtained from these fields and applies it equally to flankers on both features (with one minus this value applied to target responses). In this case, if crowding is reduced for one feature (based on a low weight for motion, for instance) then it is necessarily reduced for both. By retaining all aspects of the best-fitting models derived for the first two experiments, we can fit this model using only three free parameters (direction noise, colour noise, and late noise), making the model directly comparable to the independent models described above. Parameters were determined using the fitting approach described above, with best-fitting parameters reported in Table S3 (‘Min. weight’).

The output of these best-fitting simulations is shown in the left column of Figure S6 (square data points), plotted with conventions as in Figure 3. Results from the strong motion + strong colour condition are shown in Figure S6A, where the model successfully captures the pattern of error for colour (data points and model simulations align on the y-axis) but under-predicts the rate of error for motion (particularly for the *motion error* and *both error* conditions), where data is shifted on the x-axis. This occurs because the lower overall rate of colour crowding determines the strength of crowding for both features (i.e. because the weighting field for colour has a lower peak value, it drives performance when the minimum crowding value is taken). The model fares even worse in the weak motion + strong colour condition (Figure S6B) where the reduction in crowding for motion predicts that responses should be predominantly correct on both features. The low rate of errors in this case is driven by the large direction difference, which gives a low weight that is then applied to both motion and colour. The model similarly fails to predict sufficient errors in the strong motion + weak colour condition (Figure S6C) given the reduced weights for colour that are equally applied to motion. Altogether, the model fails to capture the errors made by observers.

**Table S3.** Best-fitting free parameter values for combined all-or-none models of crowding used to simulate the data of Experiment 3, which either take the minimum (‘Min. weight’) or maximum (‘Max. weight’) weights for crowding on both features.

| Parameter | Min. weight | Min. weight<br>(no inhibition) | Max. weight | Max. weight<br>(no inhibition) |
| --- | --- | --- | --- | --- |
| Direction noise | 0.0309 | 0.0365 | 0.0120 | 0.0192 |
| Colour noise | 0.0114 | 0.0101 | 0.0062 | 0.0090 |
| Late noise | 0.0634 | 0.0121 | 0.0948 | 0.1879 |

It is possible that part of the failure of this combined mechanism could reflect differences in the underlying populations of detectors. Most notably, the population of motion detectors involves both excitatory and inhibitory interactions whereas colour interactions are purely excitatory. We therefore set these inhibitory values to zero for both populations and associated weighting fields and re-fit the combined mechanism, again taking the minimum weight for each feature as above. Best-fitting parameters for this model are shown in Table S2, listed as ‘Min. Weight. (no inhibition)’, and the output of this model plotted against data in Figure S6, panels A-C (diamonds). The model performs similarly to the combined model with inhibition and again fails to predict the dissociable errors produced by our observers.

We next consider a combined model where flanker weights were taken as the maximum value obtained from the weighting fields for motion and colour. Here, if crowding was strong in either feature then it was strong in both. The model was otherwise identical to that described above. Beginning with a model that includes the inhibitory parameters for the motion population, best-fitting parameters are shown in Table S3, listed as ‘Max. weight’.

The output of this model is shown in Figure S6D for the strong motion + strong colour condition (squares). Here the all-or-none model does well because the strength of crowding is strong in both features, driven primarily by the higher weights for motion crowding. Interestingly, the model also performs well in the weak motion + strong colour condition (Figure S6E), matching the high degree of colour errors with a reduction in motion errors. Model outputs do however diverge in the strong motion + weak colour condition (Figure S6F), where the model predicts a high degree of errors in both features, contrary to the observed reduction in colour errors for our observers.

The ability of the model to mimic independent processes in the weak motion + strong colour condition is primarily due to differences in the stimuli used for these features. Note that model inputs for the strong crowding conditions were ±15° for direction and ±30° for hue (matching the values used for our observers). With the same weight applied to both features, the larger difference for the hue values has a greater ‘pull’ on the target response distribution (particularly given the broad tuning of detectors in these populations), increasing the chance of errors for colour. This can be seen in the strong motion + strong colour condition, where colour errors are higher than motion errors in the *both differ* condition (with both driven by the ±15° direction weights). In the weak motion + strong colour condition, the higher likelihood of these colour errors therefore pushes responses into the ‘colour errors’ quadrant, despite the overall reduction in weights (given the lower ±30° colour weights here). In contrast, the strong motion + weak colour condition is driven by higher weight values (again given the smaller difference for flanker directions than for hues), pushing errors back into the ‘both errors’ quadrant. In other words, the asymmetric performance of these maximum-probability models is driven by the different flanker values selected for motion and colour. Indeed, running these models with identical values for both features produces a more symmetric output where the model either responds with errors for both features or neither. Nonetheless, although the best-fitting parameters could mimic an independent model in some conditions, the same model necessarily fails in others.

**Figure S6.**
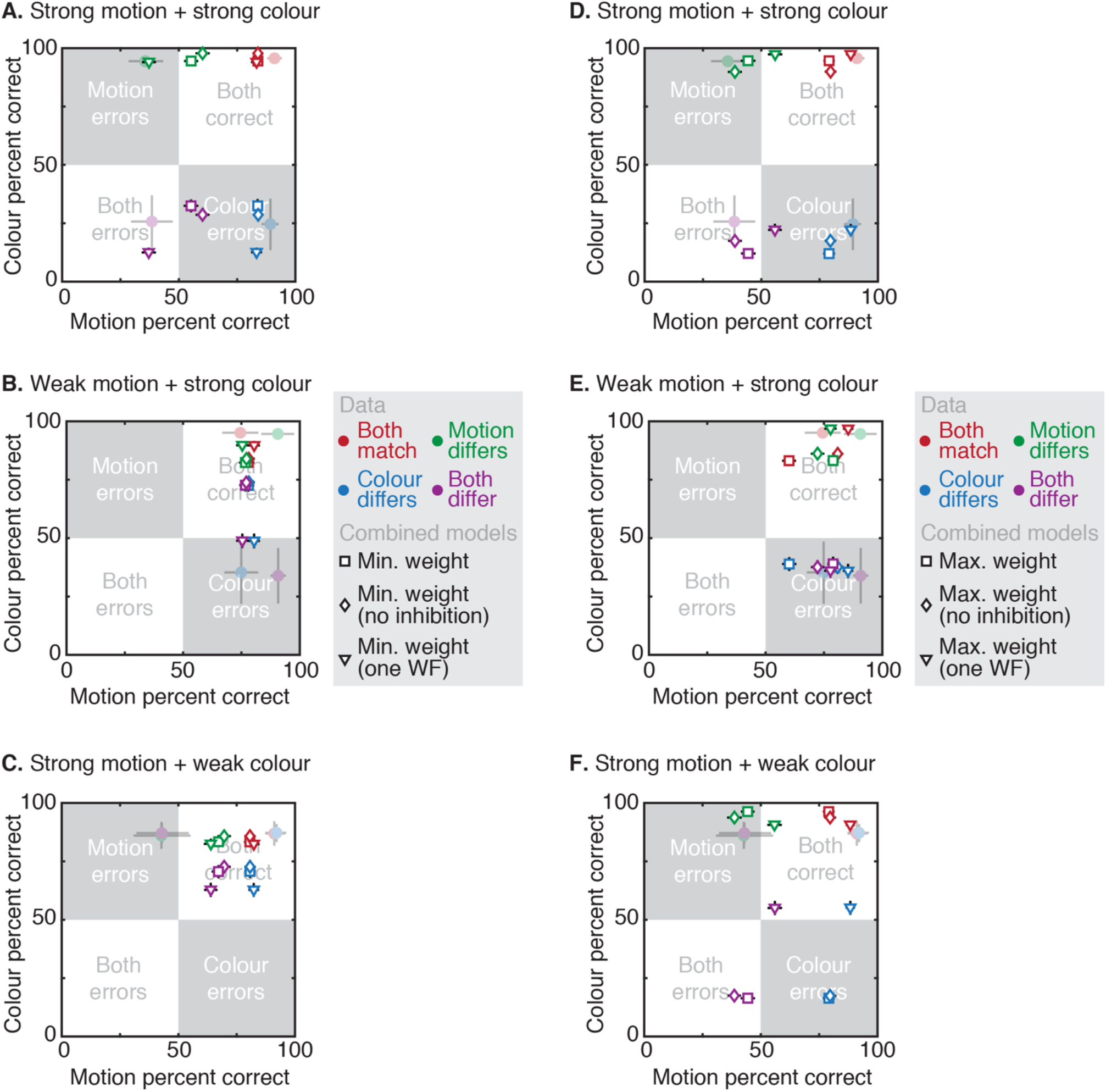
Simulations of the data in Experiment 3 by combined models of crowding. As in Figure 3, each panel plots the percent correct for the target direction on the x-axis and target hue on the y-axis. Colours show the 4 target-flanker match cases: where the 2AFC sign in each feature matches for both (red points), where the motion differs (green), the colour alone differs (blue), or both differ (purple). Data points from Experiment 3 are shown as circles with reduced opacity. Model outputs are shown for combined models taking the minimum (left panels) or maximum (right) weight across the two features. In each case models are compared with inhibitory parameters (squares), without (diamonds), or with a common weighting field (triangles). All points show the mean ±1 SEM. Panels **A.** and **D.** show data and simulations for the strong motion + strong colour crowding condition. Panels **B.** and **E.** show the weak motion + strong colour crowding condition. Panels **C.** and **F.** show the strong motion + weak colour condition.

As with the minimum-weight models above, we also re-fit this model with all inhibitory parameters set to zero. Best-fitting parameters are shown in Table S3, with outputs shown in Figure S6 (diamonds). Removing inhibition does not improve the performance of the model.

Finally, it is possible that these combined models underperformed because the weighting field for each feature dimension had distinct parameters, as carried forward from Experiments 1 and 2 (despite the same value being applied to both features on each trial). We therefore developed a model with a common weighting field, using an additional two free parameters to fit the peak and SD values of the weights to the data from Experiment 3. To simplify this approach and reduce the number of free parameters, inhibitory parameters were set to zero for these models. Even with a common weighting field, we still need to take either the minimum or maximum weight derived from the two features in order to determine whether crowding is maintained or released for both when discrepancies arise. We thus simulated both minimum- and maximum-weight models. Remaining model details were as in the other versions of this model, as was the fitting procedure. Best-fitting values of the 5 free parameters in these two models are shown in Table S4.

**Table S4.** Best-fitting free parameter values for the combined all-or-none models of crowding with a single weighting field to simulate the data of Experiment 3. Parameters are shown for one model that takes the minimum flanker weight from the weighting field for the two features and another that takes the maximum.

| Parameter | One weighting field<br>(minimum weight) | One weighting field<br>(maximum weight) |
| --- | --- | --- |
| Direction noise | 0.0321 | 0.0229 |
| Colour noise | 0.0061 | 0.0021 |
| Late noise | 0.1267 | 0.0638 |
| Weight Field SD | 103.4573 | 31.2702 |
| Weight Field Peak | 0.5223 | 0.3126 |

Consider first the minimum weight model with a common weight field, whose output is shown in the left panels of Figure S6 (triangles). In the strong motion + strong colour crowding condition (Figure S6A), the model performs well because the weights selected for both features are high. Note that percent correct performance for each feature again differs because the larger difference in the flanker hues (±30°) has a greater ‘pull’ on the target response distribution for hue than does the direction difference (±15°). This carries through to the weak motion + strong colour condition (Figure S6B), where the model comes closer to capturing the predominance of colour errors than the other minimum weight models (though still undershoots by around 15% correct). As with the other minimum-probability combined models, the predicted error rates are then vastly underpredicted in the strong motion + weak colour condition (Figure S6C) given the low weights derived from the ±150° flanker differences in colour.

The maximum-weight model similarly fails to account for observers’ performance, shown in the right panels of Figure S6 (triangles). In the strong motion + strong colour crowding condition (Figure S6D), the model performs relatively well but under-predicts the rate of motion errors given the low overall weighting field for this model (see Table S4). In the weak motion + strong colour condition (Figure S6E), the model performs as well as other maximum-probability models, given again the greater pull of the ±30° flanker differences in hue. The overall reduction in flanker weights for this model can then be seen in the strong motion + weak colour condition (Figure S6F) where overall errors are reduced to the extent that the simulated responses fall largely within the ‘both correct’ quadrant. On the whole, the model again fails to accurately capture the performance of our observers.

In order to compare these models more quantitatively, 1000 simulations were run for each of the above 2 independent and 6 combined models. In each case, simulated responses were subtracted from the mean percent correct data in Experiment 3 to obtain squared error values. Given the variation in the number of free parameters, the Akaïke Information Criterion (AIC; 88) was computed for model comparison. The resulting mean and distribution of AIC values are shown in Figure S7 for each model, where more negative values indicate better fits to the data. Although some variants of the combined model perform well in selected conditions, the independent models vastly outperform the combined models. Altogether then, both our behavioural evidence and the outcome of these simulations point to independent crowding effects that disrupt the domains of motion and colour.

MATLAB code for all of the models described in this manuscript is available at http://github.com/eccentricvision under *MotionColourCrowdModels*.

**Figure S7.**
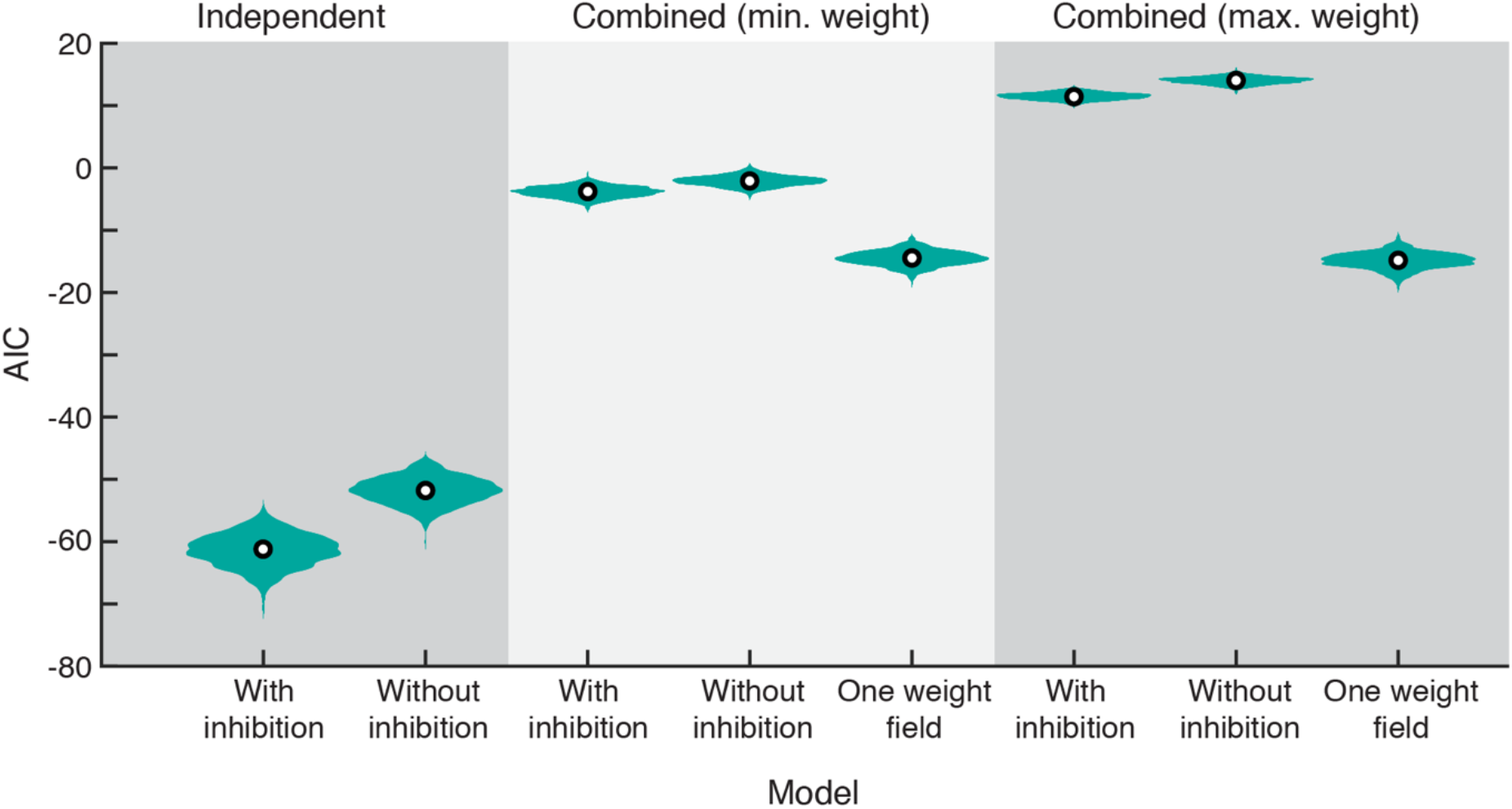
Akaike Information Criterion (AIC) values derived from the best-fitting independent and combined models of crowding. Each distribution shows the AIC value for 1000 simulations of the experiment with each model, where the width of the distribution indicates the frequency of the AIC value and circles show the mean AIC value. Two independent models are shown – one is the form shown in Figure 3 of the main text, the second excludes the inhibitory parameters in the direction-selective population. All 6 combined models show more positive AIC values, indicating worse fits. This is true for the combined models taking the minimum or maximum weight, those with or without inhibition, and those with a specifically fit weighting field.

